# Functional MRS studies of GABA and Glutamate/Glx – a systematic review and meta-analysis

**DOI:** 10.1101/2022.09.07.506899

**Authors:** Duanghathai Pasanta, Jason L. He, Talitha Ford, Georg Oeltzschner, David J. Lythgoe, Nicolaas A. Puts

## Abstract

Functional magnetic resonance spectroscopy (fMRS) can be used to investigate neurometabolic responses to external stimuli in-vivo, but findings are inconsistent. We performed a systematic review and meta-analysis on fMRS studies of the primary neurotransmitters Glutamate (Glu), Glx (Glutamate + Glutamine), and GABA. Data were extracted, grouped by metabolite, stimulus domain, and brain region, and analysed by determining standardized effect sizes. The quality of individual studies was rated. When results were analysed by metabolite type small to moderate effect sizes of 0.29-0.47 (p < 0.05) were observed for changes in Glu and Glx regardless of stimulus domain and brain region, but no significant effects were observed for GABA. Further analysis suggests that Glu, Glx and GABA responses differ by stimulus domain or task and vary depending on the time course of stimulation and data acquisition. Here, we establish effect sizes and directionality of GABA, Glu and Glx response in fMRS. This work highlights the importance of standardised reporting and minimal best practice for fMRS research.

## Introduction

### 1. Background

γ-aminobutyric acid (GABA) and glutamate (Glu), the main inhibitory and excitatory neurotransmitters in the brain, respectively, are critical for normal neurological function. GABA and Glu play an important role in perception (Edden et al., 2009; Puts et al., 2011), learning (Floyer-Lea et al., 2006), memory (Jo et al., 2014), and other behavioural functions (Paredes and Agmo, 1992; Donahue et al., 2010). GABA and Glu are known to interact, due to the fact that GABA is synthesized by using glutamic acid decarboxylase (GAD) by removing an α-carboxyl group from Glu (Cai et al., 2012). Several lines of evidence suggest that an imbalance in GABAergic and glutamatergic function is associated with neurological, neurodevelopmental, and neuropsychiatric disorders (Li et al., 2019; Tang et al., 2021; Nakahara et al., 2022). The interplay of GABA and Glu is of strong interest due to their role in excitatory and inhibitory (E/I balance) which was theorised to play important part in healthy brain function and that the disruption of E/I balance is shared by several psychiatric disorders (Yizhar et al., 2011; Ferguson and Gao, 2018).

In humans, Magnetic Resonance Spectroscopy (MRS) is the only technique that allows for the non-invasive in vivo measurement of wide range of neurometabolites including GABA and Glu (Mullins et al., 2014; Schür et al., 2016; Harris et al., 2017). MRS allows for the quantification of endogenous brain metabolites based on their chemical structure. ^1^H-proton containing metabolites each have their own distinct chemical environment and thus appear differently along a “chemical shift” axis, although with substantial overlap. Recent developments in MRS instrumental and acquisition technique have broadened our knowledge of brain neurochemistry in both clinal and research domains, and this has been extensively reviewed (Duarte et al., 2012; Faghihi et al., 2017).

While baseline GABA and Glu levels have been associated with typical and atypical brain function and behaviour (Coghlan et al., 2012; Horder et al., 2018), metabolite levels assessed at rest limit interpretation; as they cannot provide information on the temporal dynamics of GABA and Glu, which may provide insight into typical or atypical function, the relationship between GABA and Glu, task-related changes, and responses to pharmacological intervention. This has led to an increased interest in functional MRS studies, which have the potential to measure a dynamic neurochemical system.

#### 1.1 Functional MRS

Functional MRS (fMRS) refers to the use of MRS to estimate metabolite *changes* in response to external stimulation by acquiring data at different time point associated with changes in stimulus presentation. Typically, MRS spectra result from an averaged signal from repeated measurements (transients) to improve signal to noise ratio (SNR) as metabolites have an inherently low SNR due to their low concentration. A single transient refers to the data collected in each repeat (repetition time, TR) during the MRS acquisition. The often-used term ‘averages’ in MRS stems from the averaging of these transients for a single ‘average’ spectrum. Functional MRS uses the same approach but tend to measure the signal in shorter durations, or average a smaller set of transients, than in static MRS. In this study, we refer to the number of repeated acquisitions per time point as transients to avoid confusion with the act of spectral averaging. It should be noted that different acquisition sequences exist for MRS, with the most popular single-voxel MRS sequences being spin-echo point-resolved spectroscopy (PRESS) (Bottomley, 1987), stimulated echo acquisition mode (STEAM) (Frahm et al., 1989), semi localization by adiabatic selective refocusing (sLASER)(Öz and Tkáč, 2011), and spin-echo full intensity acquired localised (SPECIAL) (Kuwabara et al., 1995). Details on these approaches are beyond the scope of this work but details can be found in recent consensus work (Peek et al., 2020; Lin et al., 2021).

fMRS has been used to study wide range of brain chemistry, includes high-concentration metabolites, such as N-acetyl aspartate, creatine, and choline, to low-concentration metabolites such as lactate (see (Prichard, 1992; Chen et al., 1993; Henning, 2018; Wilson et al., 2019; Peek et al., 2020)). While the fMRS of Glu, and particularly that of GABA is of immense interest due to their critical role in brain function, fMRS of these other metabolites is not yet well-established due to technical considerations (e.g., an absence of lactate at baseline) and perhaps more difficult interpretation of its outcomes).

Glu and GABA overlap considerably with signals from glutamine (Gln) and glutathione (GSH), particularly at clinical field strength (3 T) (see 1.4). Still, despite these challenges, fMRS of GABA and Glu has been used to study neurochemical changes associated with various type of exogenous change, including pain (Gutzeit et al., 2013; Cleve et al., 2015), visual stimulation (Mangia et al., 2007; Apšvalka et al., 2015; Bednařík et al., 2015), working memory (Woodcock et al., 2018), learning and memory (Stanley et al., 2017), and motor tasks (Schaller et al., 2014; Kolasinski et al., 2019). However, substantial inconsistencies between studies exist in terms of acquisition, analysis, findings, and interpretation. To date, the body of fMRS literature on Glu and GABA has not been systematically evaluated and analysed. From hereon we refer to fMRS studies of GABA and Glu as ‘fMRS’.

#### 1.2 Limitations in estimating GABA and Glu

The measurement of GABA and Glu is challenging and contributes to variability across studies. GABA has a low concentration within the brain (1 - 2 mM), and its signal overlaps with high-concentration metabolites like NAA and creatine, as well as very similar chemical shift between Glu, Gln, and GSH. Spectral-editing techniques such as MEscher-Garwood Point-REsolved SpectroScopy (MEGA-PRESS) are often used to improve GABA resolution (Mescher et al., 1998; Edden and Barker, 2007; Near et al., 2011). These approaches rely on J-difference editing of the GABA signal, removing unwanted signal from the spectrum. For a technical review, see (Puts and Edden, 2012; Mullins et al., 2014; Wilson et al., 2019; Deelchand et al., 2021). Spectral-editing MRS techniques typically requires more transients (in the order of 8 minutes; 240+ transients for voxel sizes of 27 ml based on consensus for adequate data acquisition at 3T) compared to non-edited sequences for Glu (64 transients for voxel sizes of 8 ml at 3T) (Peek et al., 2020; Lin et al., 2021). Differences in MRS sequences, especially editing sequences, may affect the ability to interpret and reproduce studies (Terpstra et al., 2016; Baeshen et al., 2020). Whether linear-combination modelling approaches can successfully and reliably separate Glu from Gln, GABA and GSH remains inconclusive (Sanaei Nezhad et al., 2018; Zöllner et al., 2021) and thus, the composite measure Glx (= Glu + Gln) is commonly reported.

#### 1.3 Heterogeneity in fMRS approaches

There is little homogeneity regarding fMRS experimental design, stimulus type, brain region and the quality of MRS acquisition and analysis methods — all of which often depends on the research question. fMRS can typically be performed using two types of experimental paradigms, block-designs or event-related designs (Mullins, 2018). Block designs contrast metabolite measurements between acquisition blocks that are often long in duration and contain numerous stimuli and transients. Event-related approaches rely on time-locking stimulus onset with the MRS acquisition and allow for the investigation of transient metabolite levels changes immediately after stimulus onset (stimulus-locked). Block approaches typically have more SNR as more transients are averaged across per spectrum and from the summation of responses presented in close succession but have limited interpretability of stimulus-locked neurochemical responses. Effect sizes are heterogeneous, with reported observed effect sizes (*if at all* reported) range from 2% to 18% change from baseline for visual stimulation, and up to 18% change from baseline for painful stimulation (Gussew et al., 2010; Mullins, 2018; Stanley and Raz, 2018). Event-related designs are more tightly associated with stimulus timings, but often suffer from low SNR due to a limited number of transients being averaged across. Both approaches are limited by multiple unknowns such as: the response function describing the delay between stimulus and neurotransmitter change, optimal acquisition duration and timing, and optimal data analysis techniques.

#### 1.4. Our approach

One prior meta-analysis of fMRS studies focused exclusively on Glu (Mullins, 2018), however, no meta-analysis it yet to investigate the fMRS of GABA. With increasing interest in GABA and the popular concept of excitation-inhibition balance (E/I), a comprehensive meta-analysis of both GABA and Glu is of strong interest. We then further investigate potential factors that could affect outcomes in fMRS studies including fMRS design, fMRS parameters, quality of MRS studies and other source of bias.

### 2. Materials and Methods

#### 2.1 Search strategy and Inclusion criteria

A systematic search of databases (Pubmed, Ovid Medline, and Google Scholar) was performed using a search Boolean generated from *litserchr* package in R (Grames et al., 2019) combined with additional search terms based on discussion with co-author NAP (For search terms, see Supplementary Table 1). After the initial search on 21^st^ May 2021, the abstract of each article was screened to identify relevant studies using the *metagear* package in R (Lajeunesse, 2016). The studies that met the following criteria were included: 1) use of in-vivo fMRS to measure neurometabolites in the brain; 2) the study investigated changes in GABA or Glutamate (both Glu and Glx) in response to non-invasive stimuli or tasks; 3) the study participants were healthy adult humans or the study contained a healthy human control group (no psychiatric or neurological condition); 4) the study had a baseline or control condition; 5) the study was published in a peer-reviewed journal, and was written in English or translated to English via Google Translate. Relevant articles from the reference sections of included studies were identified and manually added to the analysis after being discussed with a senior author (NAP).

#### 2.2 Study selection and data extraction

Following PRISMA and PROSPERO guidelines for systematic evidence synthesis, we pre-registered this meta-analysis on Prospero (CRD42021257339) and identified relevant literature (Tricco et al., 2018). A two-stage method was used for study selection (Furlan et al., 2009). In the first stage, potentially relevant titles and abstracts were independently assessed by two investigators (DP and NAP). If the abstract was inconclusive, the full text was retrieved and assessed for eligibility. In the second stage, the investigators independently assessed the full text of potential studies selected in the first stage for their eligibility. A third investigator (JH) was consulted if disagreements persisted in both stages. Reasons for exclusion were documented.

Two investigators (DP and NAP) independently extracted the data using an identical extraction sheet. Data were extracted into four main topics of interest: 1) neurometabolite levels during fMRS; 2) study characteristics (i.e., sample size, age, gender); 3) reported MRS acquisition parameters according to the MRS-Q (e.g., MRS sequence, fMRS paradigm and timing, voxel size, TE, TR, pre- and post-processing); and 4) bibliometric data (e.g., authors, year of publication, and type of publication).

Concentrations of GABA, Glu and Glx were taken as reported by the study, mean and standard deviation (SD; mean_metab_) or as percentage change from baseline (%change_metab_). While it is possible to perform a meta-analysis on *all* data calculated as %change_metab_, the SD of these two types of data are on different scales and therefore should not be combined together (Higgins, 2011). Our approach allows data points to be combined while avoiding secondary calculation of data. The data, whether time point or time-course data, were considered as separate datapoints and compared to ‘rest’ or ‘baseline’, as long as the actual data are reported separately. Dependence of time-course data is discussed below in section 2.5. If numerical data were not explicitly reported, imputation methods recommended by the Cochrane handbook were used (Higgins et al., 2011). Data not reported in-text but in figures were extracted using WebPlotDigitizer (Rohatgi, 2021). The time from the start of the MRS acquisition to the time of metabolite measurement was also extracted. Differences between brain regions (voxels) were considered independent and therefore data from multiple brain regions acquired in a single study were extracted as independent datapoints (Peek et al., 2020). If limited studies of specific voxels were available, we grouped them based on a broader brain region (e.g., ‘frontal’, or ‘parietal’).

#### 2.3 Quality assessment

The Risk of Bias Assessment tool for Non-randomized Studies (RoBANS) (Kim et al., 2013a) was used to determine the quality of the methodological design and reporting. MRS-Q (Peek et al., 2020) and https://osf.io/8s7j9/, is a quality appraisal tool specifically designed for the systematic review of MRS studies. The MRS-Q was used to assess whether the reported acquisition methods satisfy the minimal best practice in MRS. The MRS-Q allows for assessing both the acquisition approach and whether reporting was adequate (Peek et al., 2020), and is in line with the recently published MRSinMRS (Lin et al., 2021). As the MRS-Q was designed for static MRS, its application for functional MRS experiments is discussed further in the *Discussion*). Studies were categorised into “low-quality” and “high-quality” based on the adequacy of reported MRS parameters. Studies that reported sufficient spectroscopy parameters and satisfy the consensus for adequate data acquisition were classified as ‘high quality’, studies that reported insufficient spectroscopy parameters or did not satisfy the consensus for adequate data acquisition were classified as ‘low quality’, and studies with not enough information to classify were considered ‘unsure’. While we used these terms (as per these guidelines) these do not always reflect that the study itself of low quality but perhaps did not report sufficient information per recommendation. We should also consider these in the context of history. As detailed below, we analyse data with and without inclusion of “low-quality” papers, but also perform a more dimensional approach, testing the association between effect size and acquisition parameters. Two investigators (DP and NAP) independently assessed the quality of each study using both tools. Disagreements were discussed and resolved by consensus with a third investigator (JH).

#### 2.4 Publication bias

Data were assessed for publication bias separately for each metabolite (GABA and Glu/Glx). The effect sizes were then aggregated for each metabolite within each study to avoid non-independence effects using Egger’s regression and trim-and-fill test (Duval and Tweedie, 2000; Bowden et al., 2015; Nakagawa et al., 2021). For the trim-and-fill test, a random-effects model was used on aggregated data, thus not accounting for non-independent effect sizes. Then, the Knapp and Hartung method (IntHout et al., 2014) was used to test for publication bias instead of the Wald test (Z-tests) as it has been suggested to have better performance on trim-and-fill approaches (Nakagawa et al., 2021). Aggregate effect sizes for each study were calculated by the ‘*aggregate*’ function from the *metafor* package in R (Viechtbauer, 2010). Compound symmetric structure (CS) and a conservative rho value of 0.7 were applied as per Rosenthal (1986). Data are visualized using funnel plots (Begg and Mazumdar, 1994; Sterne and Egger, 2001) with standard error (SE) as a measure of uncertainty.

#### 2.5 Data analysis

The meta-analysis was performed on the extracted data to estimate effect sizes in each study using the *Meta-Essentials* tool in R (Suurmond et al., 2017). Standardized mean differences and 95% confidence intervals (Hedge’s G) were calculated from the mean metabolite concentration change from baseline and/or the percentage change from baseline (% change), as well as through their standard deviation, allowing us to compare data reported in different units. If not specified, the first rest period was selected as baseline condition to calculate the mean difference for all fMRS designs (block, event-related and time course data).

Since data extracted from time courses are considered dependent, their effect sizes should be considered dependent as well. Therefore, time course data were first analysed separately and then sub-grouped within-study with a random variance component (Tau) weighting separately for each sub-group (Hak et al., 2016; Suurmond et al., 2017). Studies that did not allow for effect size calculation due to missing information (e.g., concentration or %change) were included in the systematic review but not in the meta-analysis. Heterogeneity of data was evaluated using I^2^ (Higgins et al., 2003). The I^2^ statistic is an estimate of proportion of variance in effect size that reflects real heterogeneity. I^2^ is a relative measure with a range from 0 to 100. Low I^2^ suggests no heterogeneity in data and no effect of moderator or potential clustering within the data. A high I^2^ suggests there are external factors and biases driving the dispersions of effect sizes, which should result in further sub-group analysis (Hak et al., 2016; Borenstein et al., 2021).

Most of the effect size estimates extracted in this current study consisted of time series data, or several datapoints came from a single study (i.e., multiple outcomes from the same participants, for example, rest versus stimulation conditions). This led to statistical dependency between measures, which can lead to errors in variance estimation of the combined effect size (Borenstein et al., 2021). To take the relationships among outcomes into account, robust variance estimation (RVE) was used. RVE has the advantage of approximating the dependence structure rather than requiring exact dependence values between effect sizes, as these are unknown for most of the studies included (Pustejovsky and Tipton, 2021). We used a conservative correlation coefficient of 0.7 for all observations (i.e. pre- and post-observations; time course data) in accordance to Rosenthal (1986)’s recommendations.

Our main aim was to identify general patterns in the fMRS responses of GABA, Glu and Glx. We then sub-grouped the data and analysed it based on type of stimulation, type of paradigm (i.e., block or event-related), and acquisition and analysis parameters (i.e., time, number of transients per time point). Beyond stimulation type we also analysed the data by brain area (region of interest). Because of variation in voxel location and limited available data for specific voxels we opted to analyse these data by region to ensure collation of data. We grouped the ROIs to optimize the number of studies yet retain a semblance of functional relevance. For example, motor cortex and medial prefrontal cortex were categorized as ‘frontal region’. We were also interested to establish whether there was an association between effect size and quality of acquisition (based on the MRS-Q). We first performed subgroup analysis on high-quality versus low-quality studies. We then estimated the correlation coefficient between effect size and number of transients and voxel size using Spearman’s rho. Finally, we explored effect size as function of time using LOESS (locally weighted least squares regression) fitting to investigate the non-linear trend of metabolite changes over the course of an acquisition, as an exploratory step to inform on potential temporal dynamics of the metabolite response (Ruppert and Wand, 1994). We do not expect this to be linear, nor do we have any a priori expectations regarding the non-linear trajectory. Only metabolite levels during stimulation periods were taken into account for this analysis; metabolite levels during breaks or periods of rest in between stimulation periods were excluded. The start of MRS acquisition was considered as t = 0 s.

### 3. Results

#### 3.1 Study selection

The initial search returned 3,385 studies. After automatic removal of duplicates, 3,383 studies were eligible for abstract screening. 3,329 studies were excluded in the abstract screening stage for the following reasons: additional duplicate studies (n = 538); irrelevant topic (n = 2,778); and animal studies (n = 13). This resulted in 54 studies eligible for full-text screening, resulting in an additional four studies excluded due to insufficient detail, and one study excluded due to it being a meta-analysis. Finally, a total of 49 studies were included in this study. A PRISMA flow diagram can be found in Figure 1.

**Figure 1:**
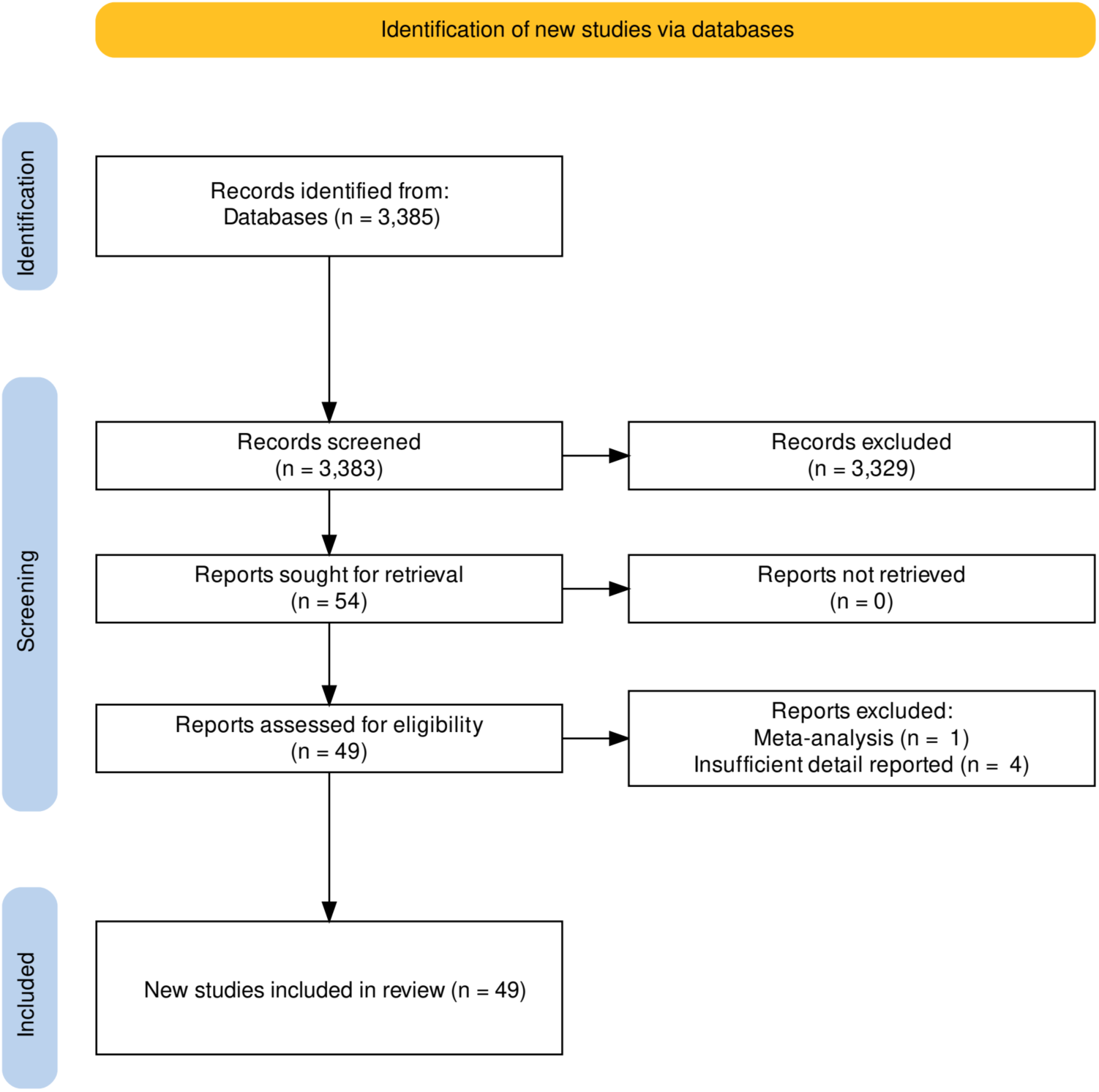
PRISMA flow diagram (Page et al., 2021; Haddaway et al., 2022)

#### 3.2 Study characteristics

##### 3.2.1 Spectroscopy

Thirty-one of the fMRS studies were performed on 3 T MR-systems, 15 at 7 T, two studies were performed at 4 T, and one study at 1.5 T. The most commonly (18 studies) used non-spectral-editing sequence was PRESS (Bottomley, 1984; Klose, 2008) six studies used STEAM, and five studies used sLASER. For spectral-editing sequences, 10 studies used MEGA-PRESS, six used SPECIAL, two studies used MEGA-sLASER, and one study used each of BASING or STRESS. Two studies reported the use of more than one editing sequence (Table 1). To measure fMRS GABA, 10 studies used MEGA-PRESS, three studies used SPECIAL, two studies used MEGA-sLASER, two studies used MEGA-sLASER, two studies used sLASER and one study used STEAM. To measured Glu and Glx, 18 studies used PRESS, 10 studies used MEGA-PRESS, six studies used STEAM, six studies used SPECIAL, five studies used sLASER, one study used each of BASING or STRESS.

**Table 1.**
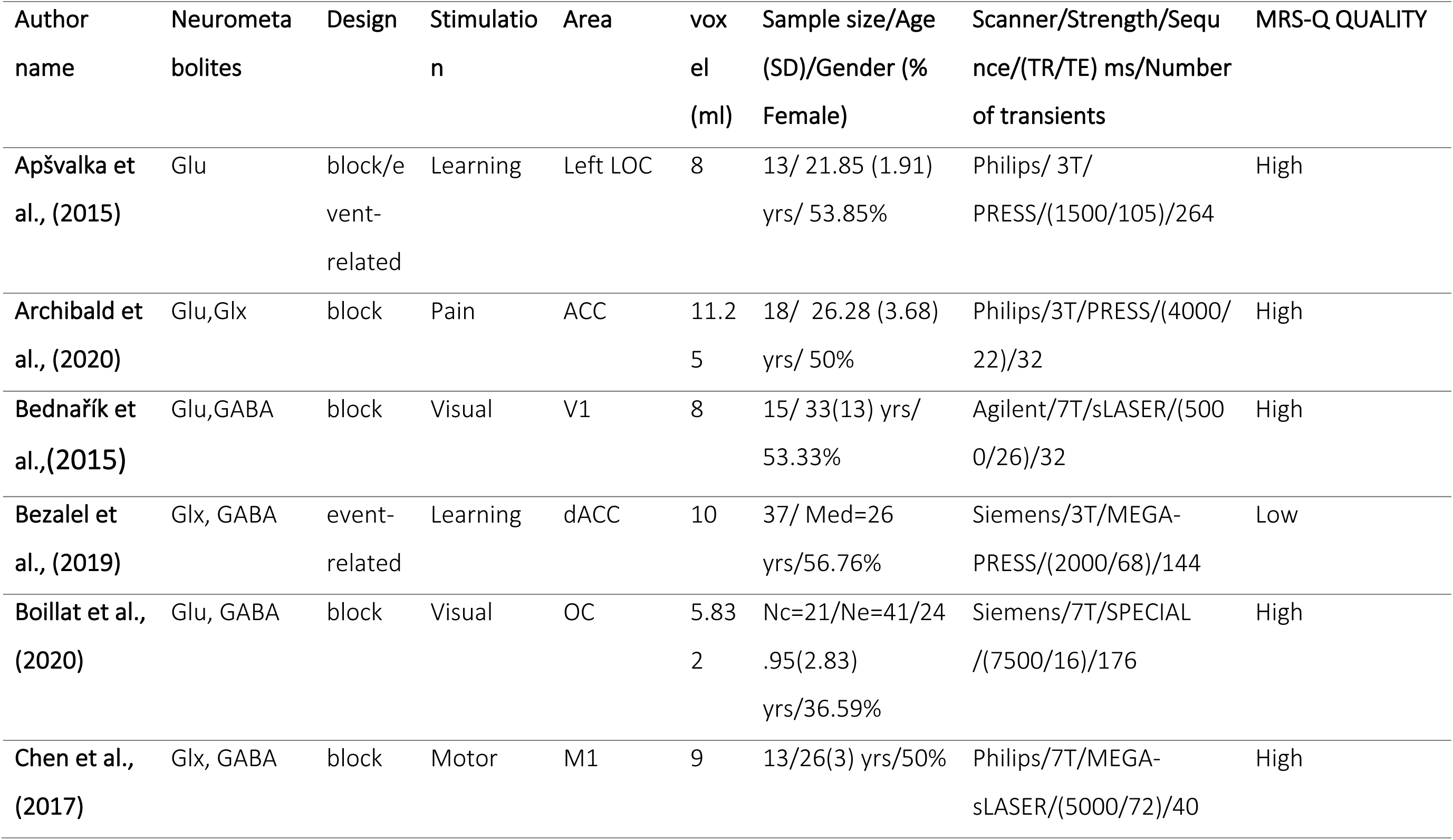
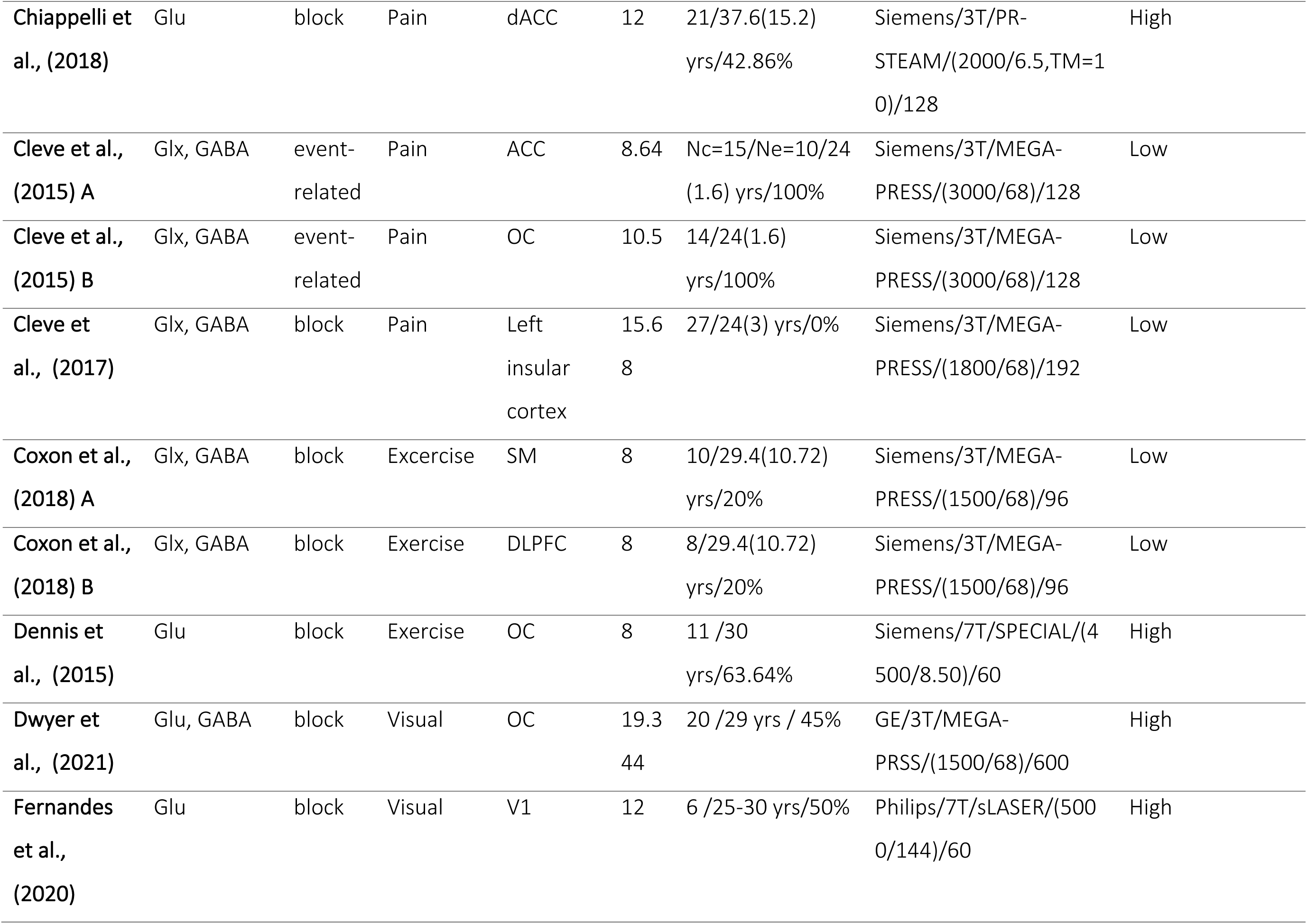
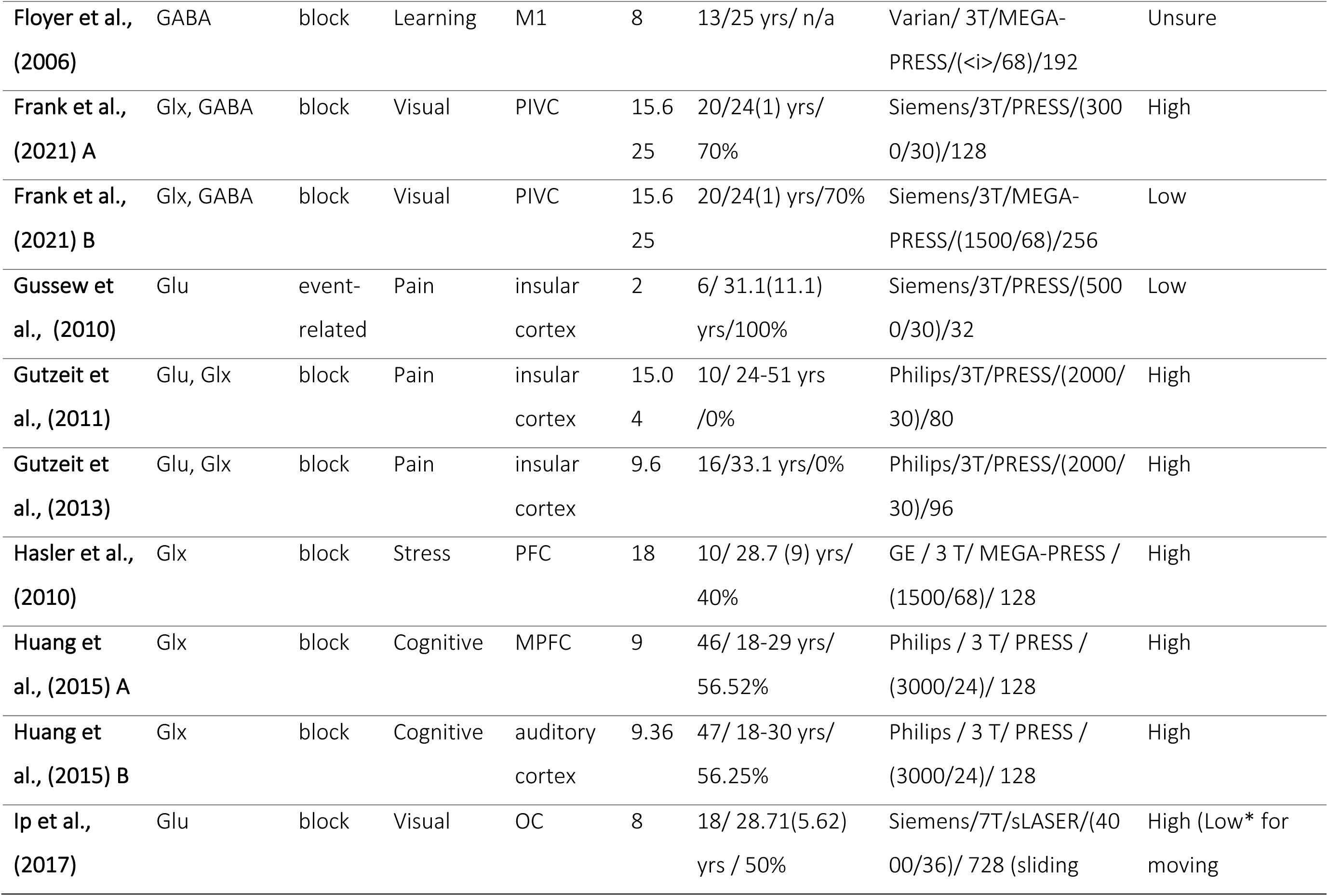
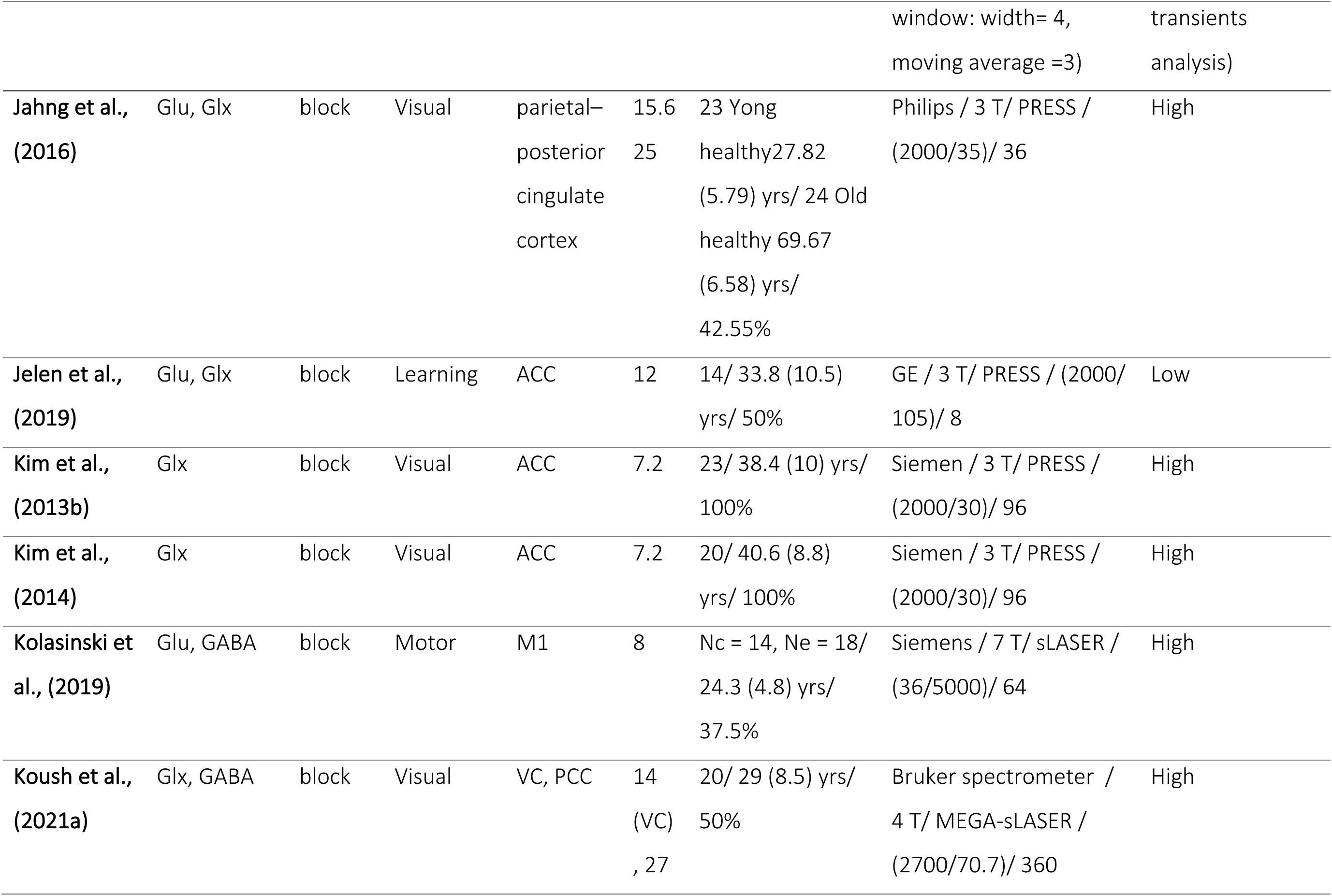
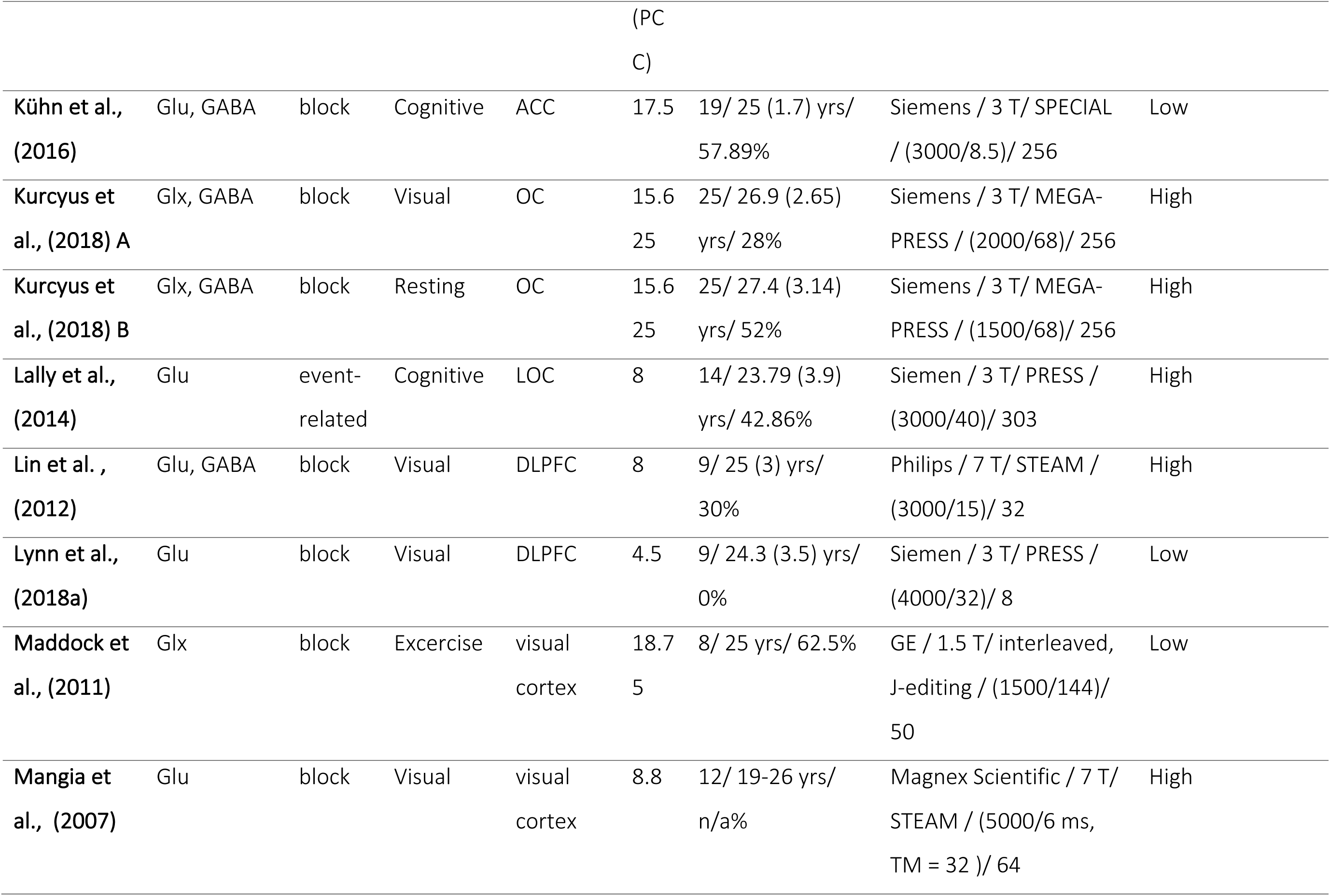
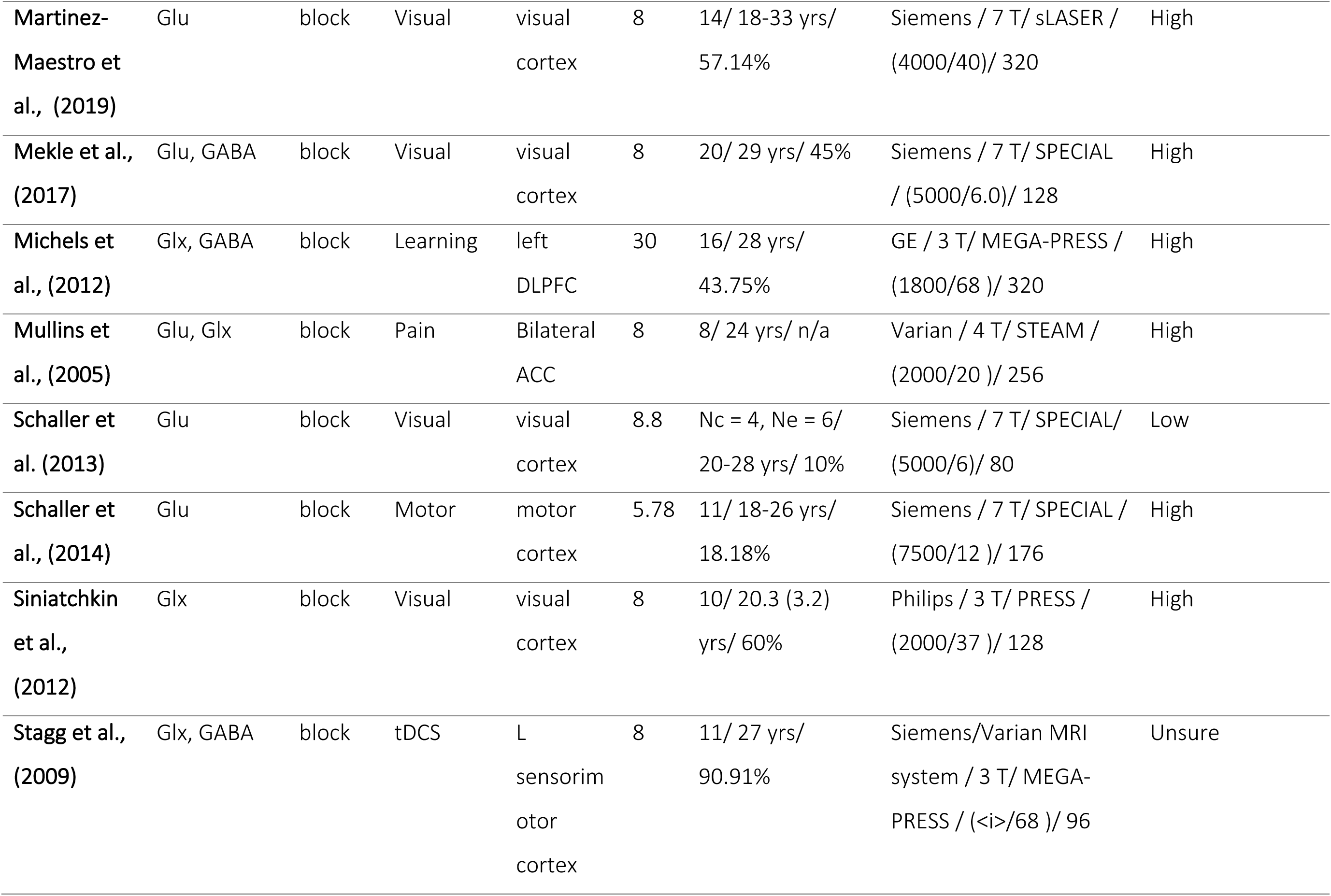
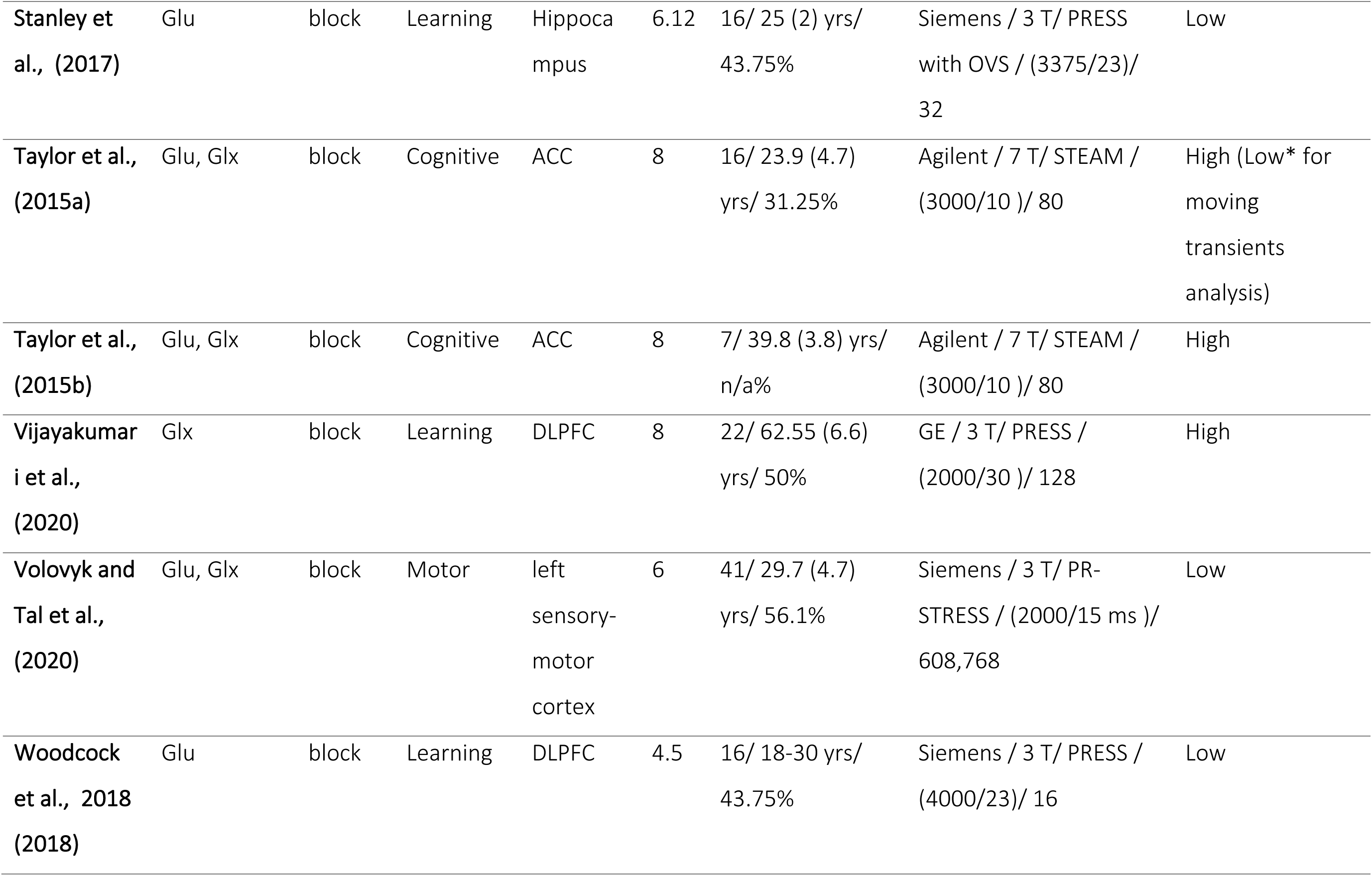
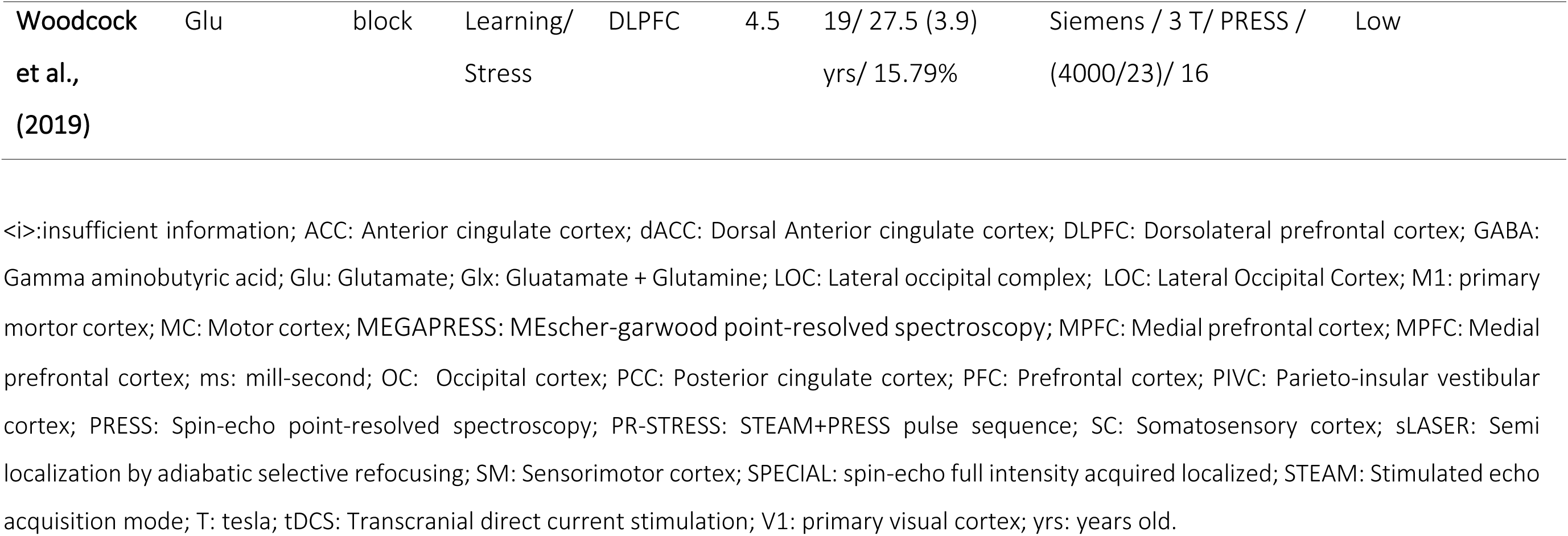
Studies characteristic

##### 3.2.2 Neurometabolites

Fifteen studies investigated only Glu levels, seven studies investigated only Glx, nine studies reported both Glu and Glx levels. Seven studies investigated both Glu and GABA, while ten studies investigated both Glx and GABA, and one study reported only GABA. See Table 1 for details.

##### 3.2.3 Stimulus domains and brain regions

We grouped studies into 8 stimulus domain categories. These domains were visual (n = 20), pain (n = 8), learning (n = 7), cognition (n = 5), motor (n = 4), stress (n = 2), tDCs (n = 1), and exercise (n = 3). Studies were considered to fall into the visual domain if they contained visual stimulation (i.e., flashing checker board, rotating checker board, visual attention tracking, and video clips) the pain domain if they contained stimulus that elicit pain (i.e., heat pain, dental pain and electric shock) learning domain if they contained learning paradigm (i.e., object recognition, reinforcement learning, n-back task (for short-term memory/implicit learning and working memory), cognition if they contained cognitive task (i.e., Stroop task, imaginary swimming and categorization of either object or abstract stimuli), motor if they contained motor response (i.e., hand clenching and finger tapping), stress if they contained psychological stress, and pharmacological stress and exercise if they contained measurement of evaluation of heart rate to exercise.

The studies were grouped in six different brain regions of interest (ROI). The most studied ROI was the occipital ROI for Glu/Glx and GABA. Additional details of MR-parameters and fMRS experiment designs are presented in Table 1. Figures 2A and 2B summarise studies by brain ROIs investigated for Glx/Glu and GABA, respectively, and additionally reports on stimulus domain.

**Figure 2:**
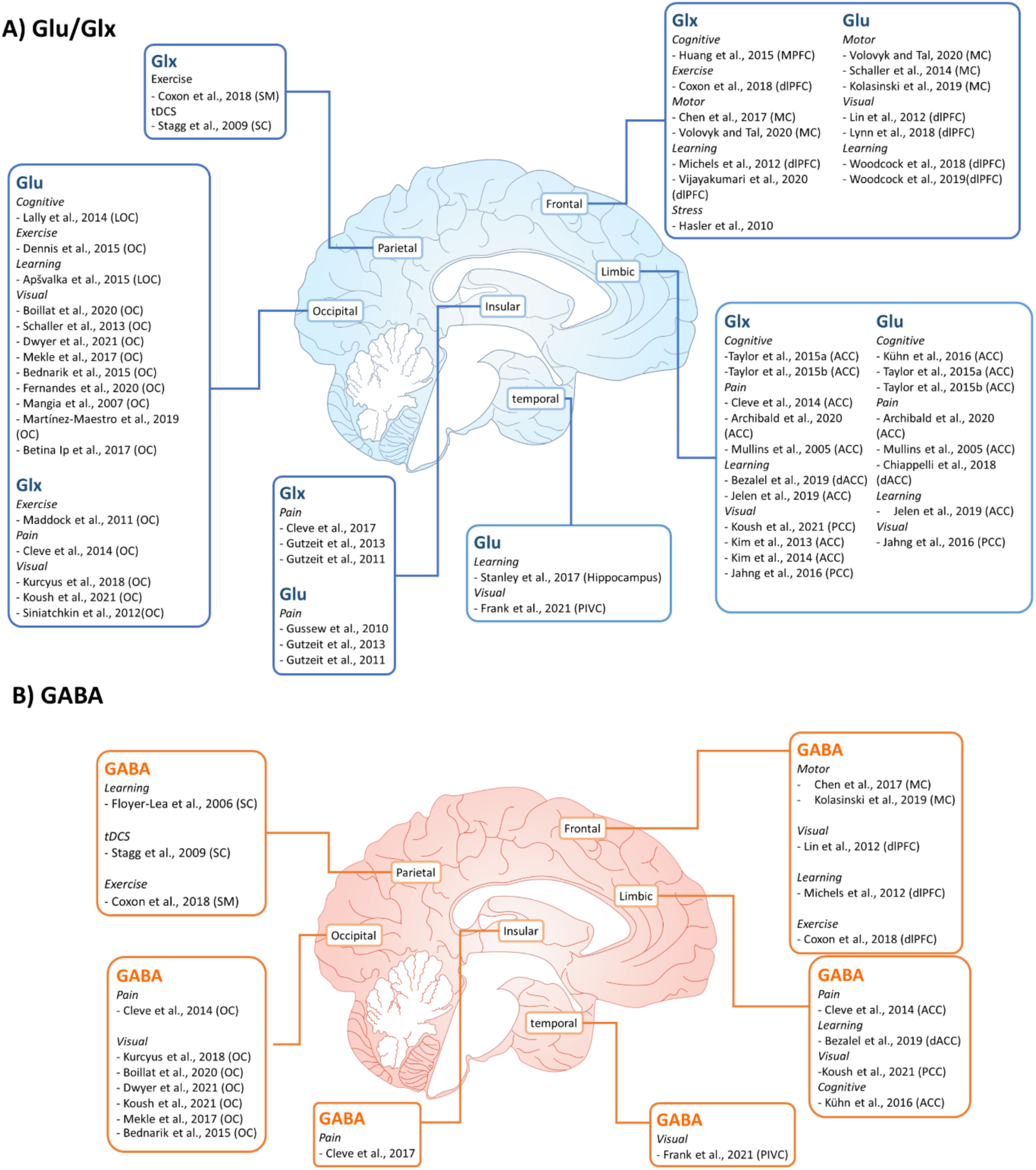
(A) Brain ROIs and stimulus domains of included fMRS studies of Glu/Glx. (B) Brain ROIs and stimulus domains of included fMRS studies of GABA. Note that brain ROIs were generalized by the authors to optimize inclusion.

#### 3.3 Quality assessment

##### 3.3.1 MRS-Q

Most studies (n = 31/49, 63.3%) satisfied the MRS-Q criteria of standardized reporting and best practice and were assessed to be of high quality (Figure 3A). Eighteen studies (36.7%) were assessed as low quality due to inadequate MRS parameters according to MRS-Q, mostly due to an insufficient number of transients or small voxel sizes (see Discussion for further consideration of using baseline MRS quality assurance approaches for fMRS). Among these low-quality studies, nine used spectral-edited fMRS(Maddock et al., 2011; Schaller et al., 2013; Cleve et al., 2015, 2017; Kühn et al., 2016; Coxon et al., 2018; Bezalel et al., 2019; Volovyk and Tal, 2020; Frank et al., 2021), while eight were non-edited (Gussew et al., 2010; Taylor et al., 2015a; Betina Ip et al., 2017; Stanley et al., 2017; Lynn et al., 2018a; Woodcock et al., 2018, 2019; Jelen et al., 2019). For high quality studies, nine studies used spectral-edited fMRS (Hasler et al., 2010; Michels et al., 2012; Schaller et al., 2014; Dennis et al., 2015, 2015; Chen et al., 2017; Mekle et al., 2017; Kurcyus et al., 2018; Boillat et al., 2020; Dwyer et al., 2021) and 24 were non-edited (Mullins et al., 2005; Mangia et al., 2007; Gutzeit et al., 2011, 2013; Lin et al., 2012; Siniatchkin et al., 2012; Kim et al., 2013b, 2014; Lally et al., 2014; Apšvalka et al., 2015; Bednařík et al., 2015; Huang et al., 2015; Taylor et al., 2015b, 2015a; Jahng et al., 2016; Betina Ip et al., 2017; Chiappelli et al., 2018; Kolasinski et al., 2019; Martínez-Maestro et al., 2019; Archibald et al., 2020; Fernandes et al., 2020; Vijayakumari et al., 2020; Frank et al., 2021; Koush et al., 2021b). Two edited-fMRS studies (Floyer-Lea et al., 2006; Stagg et al., 2009) reported insufficient information regarding the MRS parameters and were identified as ‘unsure’.

**Figure 3:**
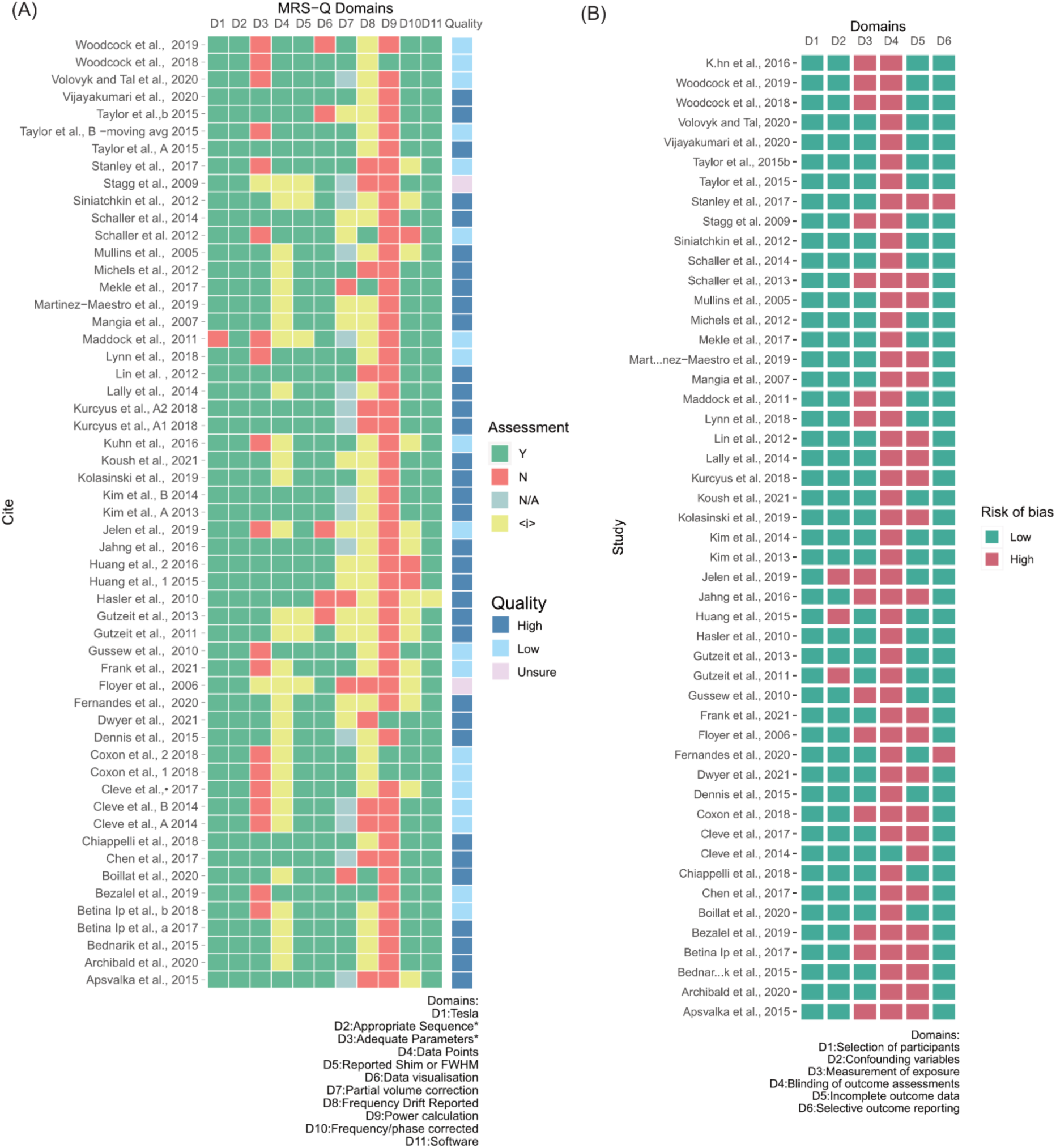
(A) MRS-Q assessment of MRS studies (B) Risk of Bias Assessment Tool for Nonrandomized Studies (RoBANS) quality.

##### 3.3.2 RoBANS

The risks of biases assessed using the RoBANS are summarized in Figure 3B. According to the RoBANS assessment, all but one study was considered to have a high risk of bias due to non-blinding of outcome, primarily due to participants or experimenters being aware of receiving/delivering a functional paradigm. Only one study explicitly reported blinding of outcome. Given the nature of fMRS experiment as a pre-post intervention study, some fMRS experiment designs might be impossible to blind. While there may be potential bias due to the fMRS examiner or participant being aware of the stimulus being given, the order of stimuli is often unknown to participant and therefore ‘blind’ to the stimulus paradigm. Yet, this bias needs to be considered as it may impact the results (e.g., participant may behave differently when the purpose is known, experiments may bias their analysis based on the paradigm). Blinding criteria are likely more relevant for pharmacological studies than for typical fMRS experiments.

fMRS studies are often required to exclude data with unsatisfactory spectral quality. While this is common in MRS, based on the RoBANS criteria, studies with incomplete outcome data would be identified as high risk. Given above criteria, 55.1% of studies were considered high-risk. Twenty-two studies (44.9%) stated that all data were included. Two studies (4.1%) were of high risk of bias for selective outcome reporting as they did not fully report all available outcomes. Bias of inadequate measurement was also identified via the MRS-Q by assessing whether studies reported adequate MRS parameters; 70% of all studies included were assessed to be at low risk of bias in this domain. No study reported potential bias in selection of participants.

#### 3.4 Publication bias

The summary for the Egger’s and Trim-and-fill test for publication bias are showed in Table 2.

**Table 2.**
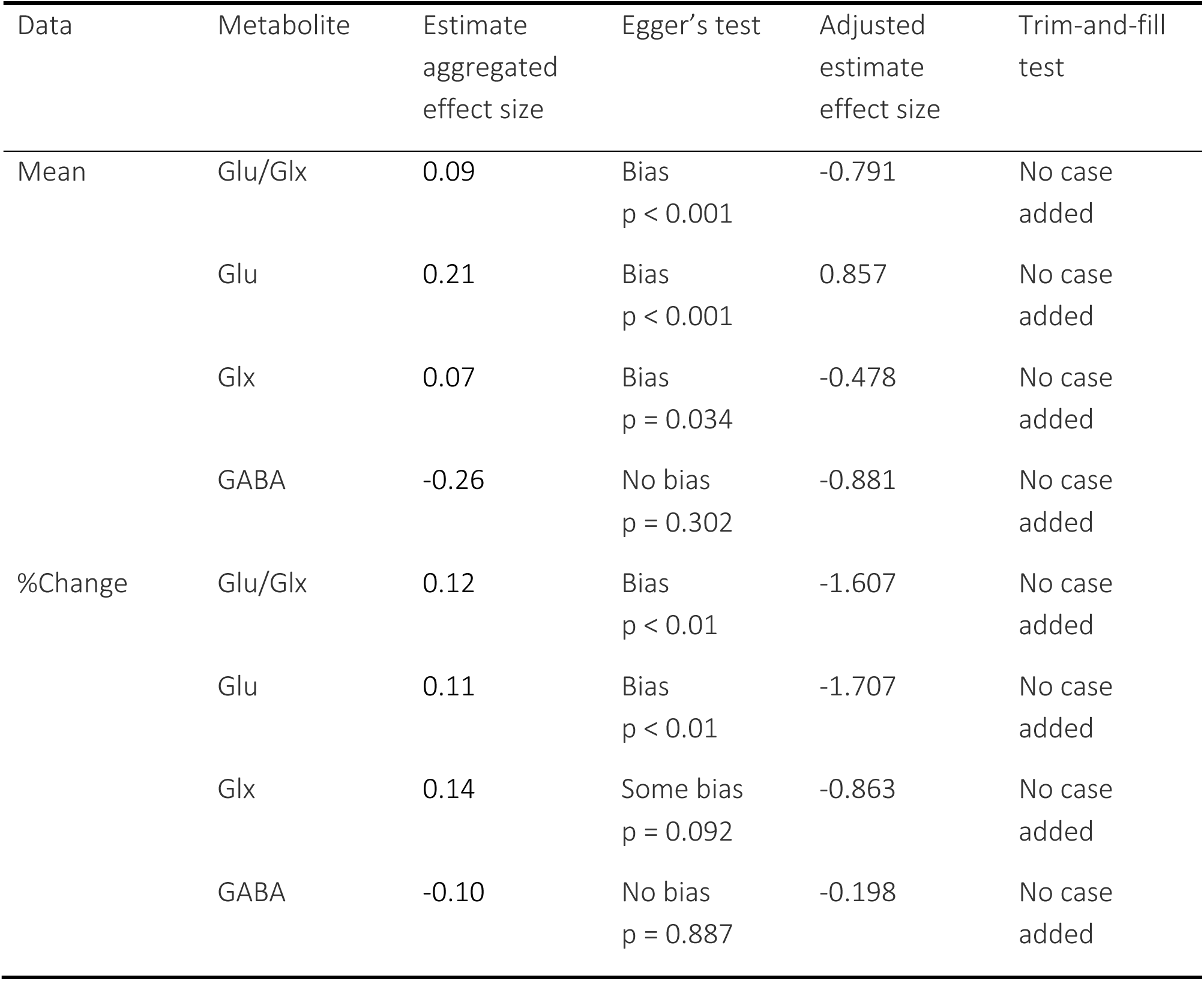
A summary of the publication bias results from both Egger’s test and Trim-and-fill test

##### 3.4.1 Egger’s regression test and funnel plot

Egger’s regression test (Egger et al., 1997) is a quantitative asymmetry test based on a simple regression model. The funnel plot illustrates the effect size of each study on the x-axis and standard error on the y-axis, without the publication bias the studies should roughly followed the funnel shape with symmetric distribution of datapoints (Lin and Chu, 2018). No asymmetry in small and large effect sizes was found for GABA (both %change_GABA_ and mean_GABA_). However, Egger’s test suggested significant asymmetry (p < 0.05) for Glu/Glx, as well as for Glu and Glx when analyzed separately, except for %change_Glx_. Supplementary Table 2 shows the results from the Egger’s regression test including the estimated effect sizes adjusted for publication bias. Supplementary Figure 1 shows the funnel plot using SE as a measure of uncertainty, color coded by stimulus domain. These data suggest that studies of Glu/Glx were asymmetrical due to an absence of small effect size positive direction studies.

##### 3.4.2 Trim-and-fill

The trim-and-fill method is a non-parametric test that was used to visualize and correct data asymmetry due to publication bias (Duval and Tweedie, 2000). The principle of the method is to ‘trim’ the studies with publication bias causing plot asymmetry, and to use the trimmed funnel plot to estimate the estimated the true centre of the funnel plot, then ‘filling’ or added the trimmed studies and their missing counterpart studies (not reported due to publication bias). Based on the method, no study was added via the trim-and-fill test; therefore, the estimated effect sizes remained the same. All data demonstrated moderate to high heterogeneity with I^2^ values of 60% - 90% (Supplementary Table 3). This means that the variability and inconsistency across study are from the true heterogeneity in the data and not by chance (Higgins et al., 2003). Trim-and-fill analysis suggested there were no potential missing studies due to bias (Supplementary Figure 2). Due to presence of between-study heterogeneity in this current study, the interpretation of these results needs to be treated with care (Terrin et al., 2003; Ioannidis and Trikalinos, 2007; Shi and Lin, 2019).

#### 3.5 Meta-analysis

##### 3.5.1 Effect of fMRS-design

###### Neurometabolite levels across all studies

When we considered change in metabolite levels across studies regardless of stimulus domain, brain ROI, or other factors (e.g., voxel size, number of transients), mean_Glu_ and mean_Glx_ increased significantly compared to the respective baseline condition (Hedge’s G_Glu_mean_ = 0.37, 95% CI: 0.09 – 0.645, I^2^ = 86.83 and G_Glx_mean_ = 0.29, 95% CI: 0.035 – 0.555, I^2^ = 87.71 respectively). The *percentage* change between baseline and active conditions in Glu was positive on average (Hedge’s G_Glu_pct_ = 0.47, 95% CI: 0.158 – 0.789, I^2^ = 82.81). No significant change was observed for GABA studies for either mean or percentage change when compared to baseline (Figure 4A).

**Figure 4:**
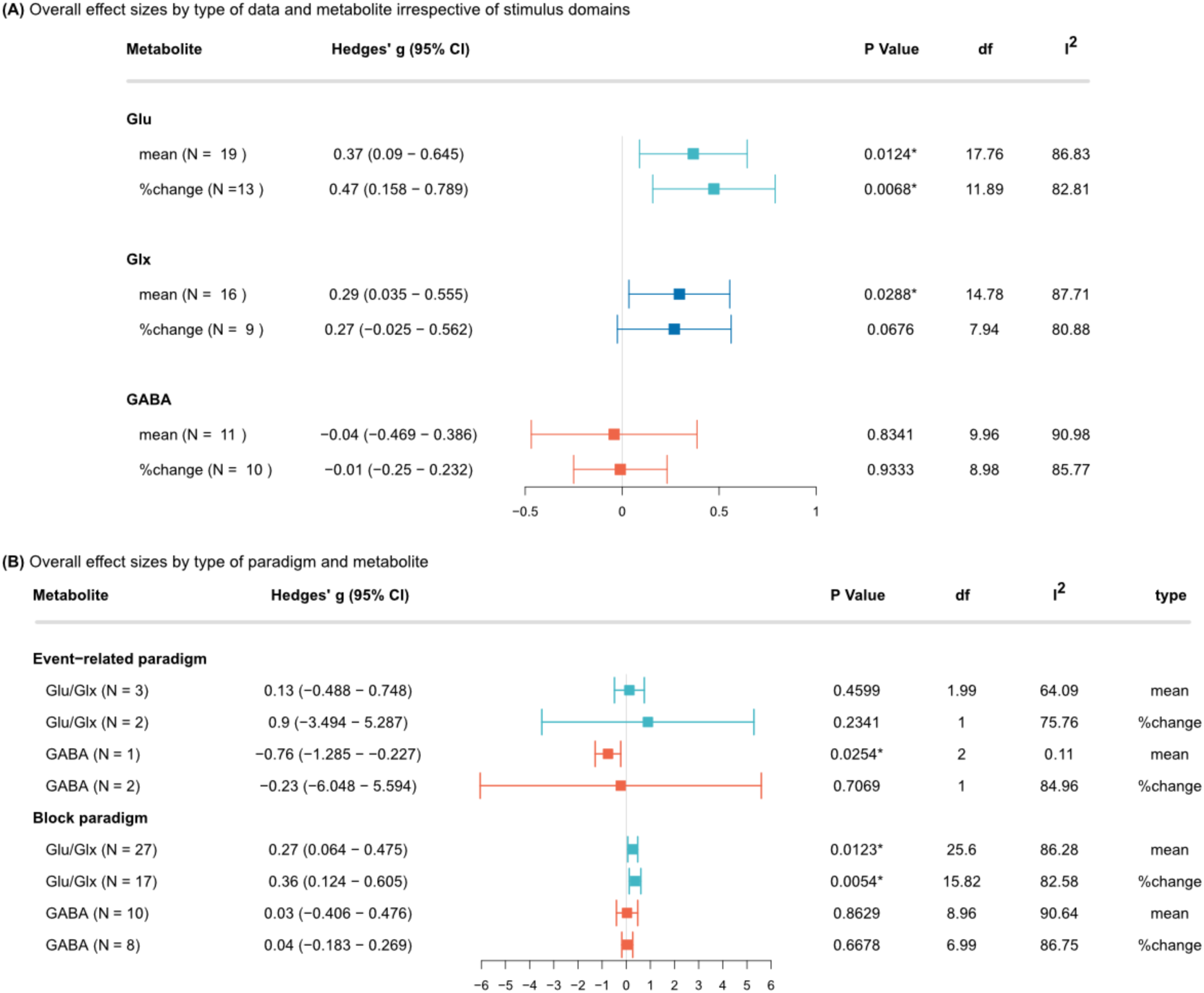
(A) Overall effect sizes by type of data and metabolite irrespective of stimulus domains. (B) Overall effect sizes by type of paradigm and metabolite. N: number of studies included; Glu/Glx: Glu or Glx studies; I^2^: I^2^ index for heterogeneity. A high I^2^ suggests there are external factors and biases driving the dispersion of effect sizes. *Statistically significant at p < 0.05, and at p < 0.01 when the degrees of freedom < 4 for RVE t-tests.

###### Neurometabolite levels by type of paradigm

When effect sizes were computed by type of paradigm regardless of brain ROI and stimulus domain, block designs showed lower confidence intervals in effect size relative to event-related designs and a significant overall positive change in Glu/Glx for both mean and %change (Hedge’s G_Glu/Glx-mean_ = 0.27, 95% CI: 0.064 – 0.475, I^2^ = 86.28; Hedge’s G_Glu/Glx-%change_ = 0.36, 95% CI: 0.124 – 0.605, I^2^ = 86.28) (Figure 4B). A significant reduction in mean GABA was observed for event-related designs (Hedge’s G_GABA_ = −0.76, 95% CI: −1.285 – −0.227, I^2^ = 0.11), but no significant change was observed for block paradigms. It must be noted that the significant effect observed here is of one study only, thus the interpretation of the result must be treated with care.

##### 3.5.2 Neurometabolite levels by stimulus domains

All stimulus domains that demonstrated a significant change from baseline contained only one individual study with 3 to 9 within-study outcomes (i.e., were driven by single studies that had multiple results at different timepoint, metabolite changes as a function of time or different types of stimuli within a single study). The percentage in GABA level increased positively during exercise (Hedge’s G_GABA-mean_ = 0.46, 95% CI: 0.023 – 0.906, I^2^ = 0.7). On the other hand, the %change_Glu/Glx_ was positive during learning (Hedge’s G_Glu/Glx-%change_ = 0.29, 95% CI: 0.106 – 0.469, I^2^ =0.23), mean GABA showed negative change from baseline (Hedge’s G_GABA-mean_ = −0.76, 95% CI: −1.285 – −0.227, I^2^ =0.11) during learning. Mean GABA and %change in GABA showed significant change in the opposite direction in the motor domain (Hedge’s G_GABA-mean_ = −0.76, 95% CI: −1.485 – −0.044, I^2^ = 0.6; Hedge’s G_GABA-%change_ = 0.32, 95% CI: 0.184 – 0.459, I^2^ = 0). Stress stimulation was associated with a significant negative change for GABA (Hedge’s G_GABA-mean_ = −0.87, 95% CI: −1.609 – −0.129, I^2^ = 0.69). During transcranial direct current stimulation, GABA showed a negative %change (Hedge’s G_GABA-%change_ = −0.12, 95% CI: −0.238 – −0.006, I^2^ = 0). There were no significant changes related to visual stimulation for any measure of Glu/Glx and GABA (Figure 5). Again, it must be highlighted that only 1-2 studies were included in these results with statistical significance, thus these findings need to be interpreted cautiously.

**Figure 5:**
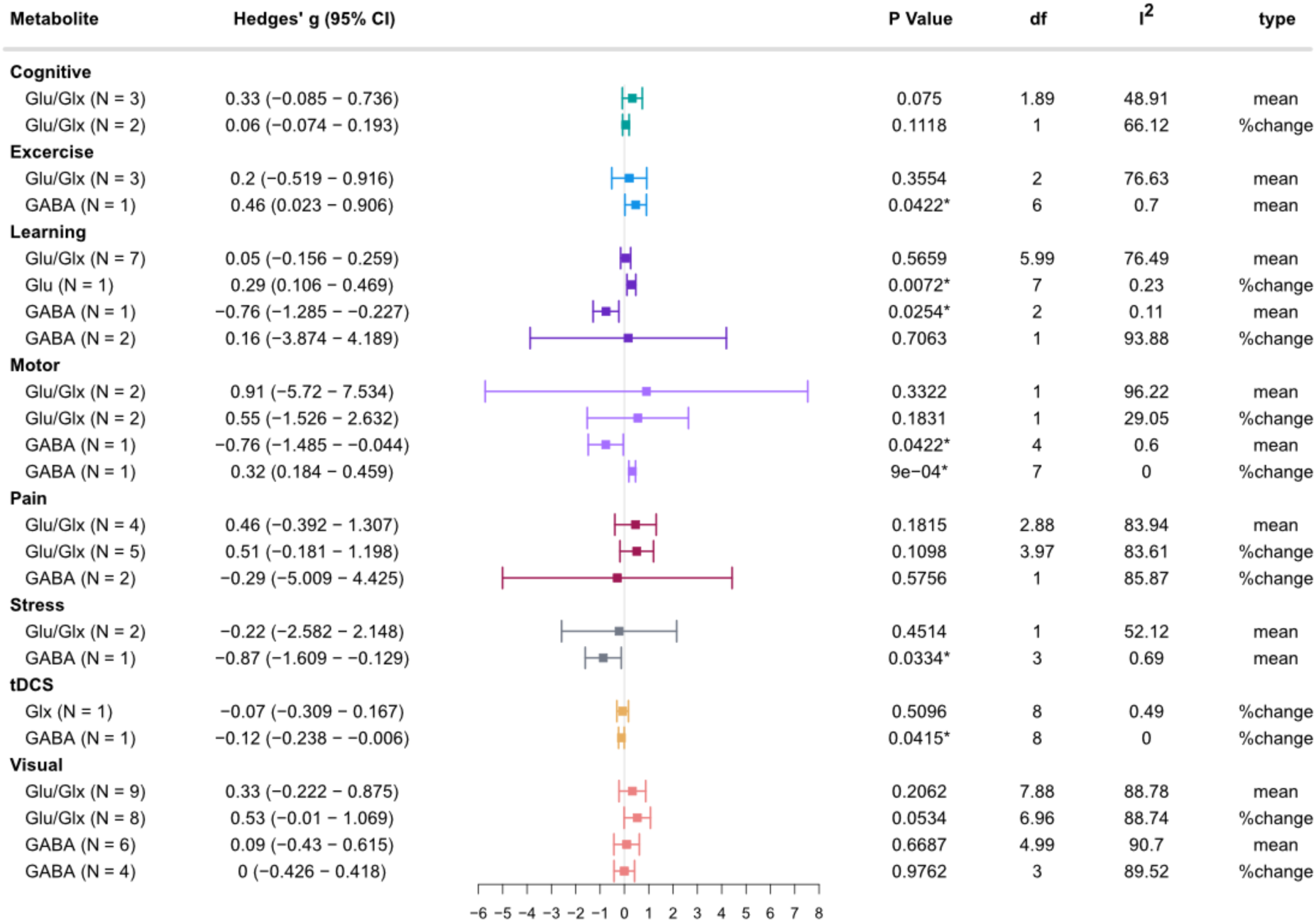
Overall effect sizes by type of stimulus and metabolite. N: number of studies included; Glu/Glx: Glu or Glx studies; I^2^: I^2^ index for heterogeneity. A high I^2^ suggests there are external factors and biases driving the dispersions of effect sizes. * Statistically significant at p <0.05, and at p <0.01 when the degrees of freedom < 4 for RVE t-tests.

##### 3.5.3 Neurometabolite levels by ROI studied

When we investigated the neurometabolites by ROI, only a few studies were included for each metabolite. Across neurometabolites, regardless of stimulus domain, every ROI except for the limbic ROI showed a significant difference in neurometabolite levels compared to the baseline condition. The occipital ROI comprised most of the studies included (n = 22 across metabolites). Pooled effect sizes from six studies in occipital ROIs observed an overall increase by %change of Glu/Glx (Hedge’s G_Glu/Glx-%change_ = 0.84, 95% CI: 0.089 - 1.588, I^2^ = 88.73). This was surprising since stimulation in the visual domain themselves showed no significant effect. This may be because the effect of visual stimulation was not only tested in visual cortex but across different ROIs (see Figure 2 and Table 1). Significant increases compared to the baseline condition were also observed for frontal %change_GABA_ (Hedge’s G_GABA-%change_ = 0.35, 95% CI: 0.046 – 0.649, I^2^ =80.86) and insular mean_Glu/Glx_ level (Hedge’s G_Glu/Glx-mean_ = 0.52, 95% CI: 0.094 – 0.95, I^2^ = 75.97). Due to limited available data, temporal and parietal ROI only had one study included for each analysis, except for percentage change in parietal GABA. While significant differences were observed, these data show very low heterogeneity (I^2^ = 0 – 0.23). This might suggest a potential bias in over- or under-estimating the observed effects since these results are from within-study outcomes. Data by ROIs are shown in Figure 6.

**Figure 6:**
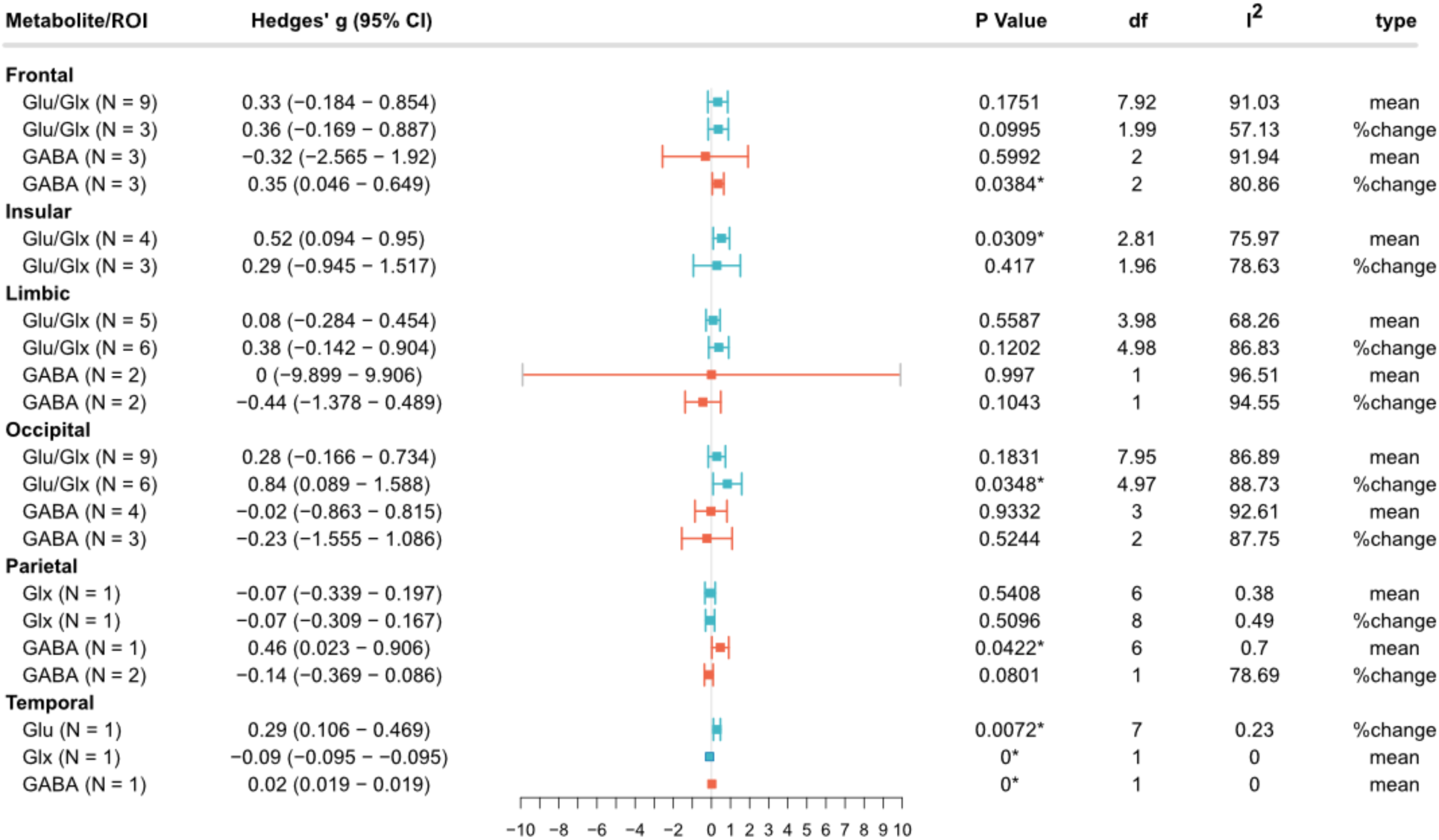
Overall effect sizes by ROIs. Glu/Glx: Glu or Glx studies; N: number of studies included; I^2^: I^2^ index for heterogeneity. A high I^2^ suggests there are external factors and biases driving the dispersions of effect sizes. * Statistically significant at p < 0.05, and at p < 0.01 when the degrees of freedom < 4 for RVE t-tests.

##### 3.5.4 Effect sizes in relation to time

Several studies had time-course data available, and we were therefore able to explore effect sizes based on ‘time-in-acquisition’ (see Figure 7). The results show different temporal fluctuation for GABA/Glu/Glx in different stimulus domains. The fitted line (LOESS) suggests potential metabolic response patterns; GABA tends to start high but then decreases with increasing time-in-acquisition in learning paradigms. For Glu/Glx_mean_, three studies were included for exercise stimulus and one study was included for each of visual, learning and stress. For mean_GABA_, one study was included for visual stimulus. For %change_Glu/Glx_, four studies were included for visual stimulus, two studies for learning, and one study each for motor and cognitive. For %changeGABA, one study was included for motor stimulus. The %change_Glu/Glx_ tends to increase with increasing stimulation for visual paradigms only. There were no clear patterns for mean_GABA_ and mean_Glu/Glx_. It should be noted that while this is interesting, the amount of available data included is too small to make a firm conclusion.

**Figure 7:**
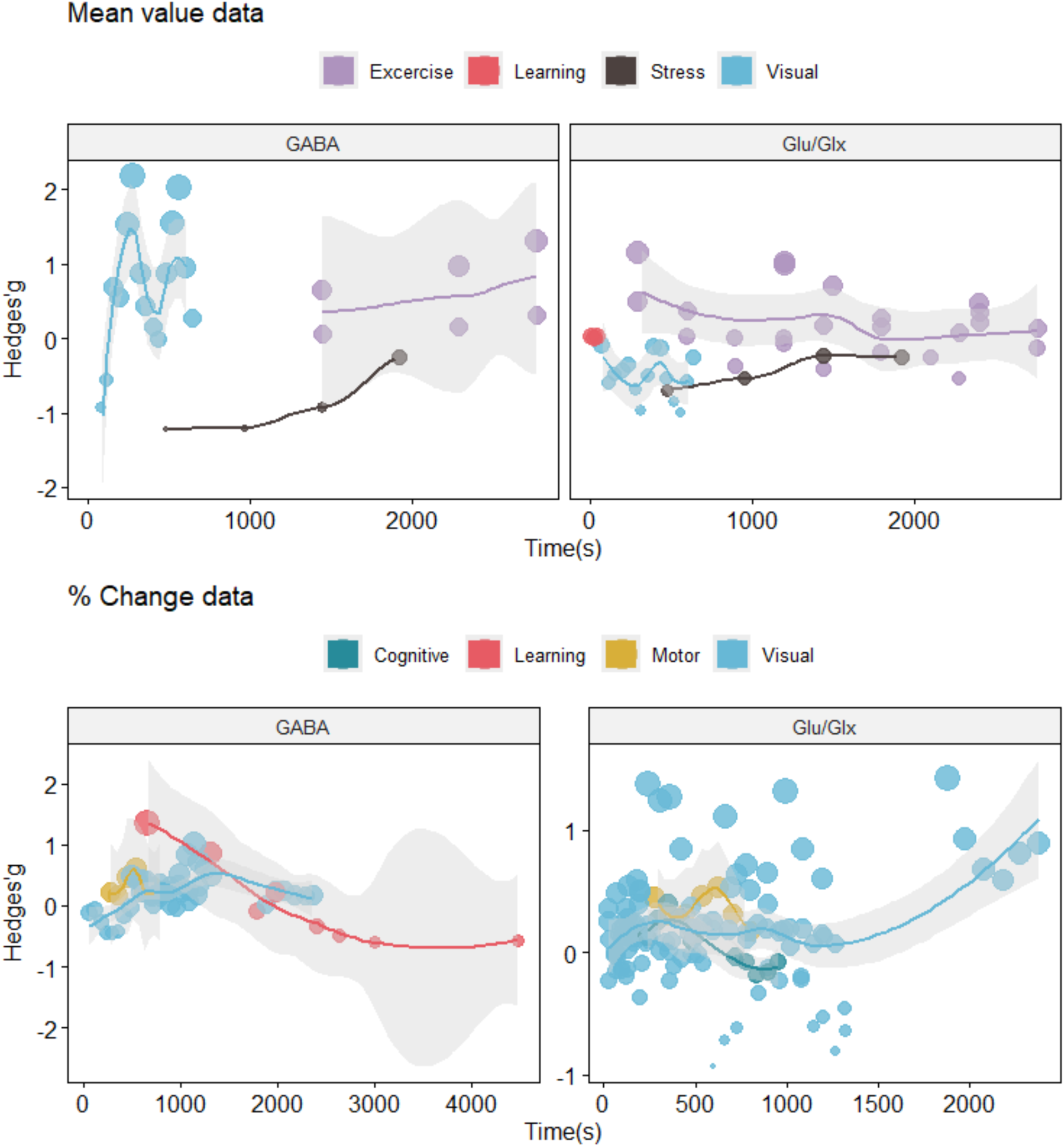
Effect size of each study in relation to time of data acquisition during fMRS. Only metabolite levels during stimulus periods were included. Time 0 s is considered the start of the MRS acquisition. The size of dots represents the weight of the effect size.

#### 3.6 Effect of fMRS-parameters

##### 3.6.1 Effect of quality based on the MRS-Q

Supplementary Figure 3 illustrates data when only ‘high quality’ studies were included. Generally, Glu/Glx show a positive trend while GABA shows a small negative trend for mean_GABA_ compared to baseline. These findings are in agreement with section 3.5.1 where we did not consider study quality. Unlike in section 3.5.1, however, the change in mean_Glu/Glx_ was not significant from baseline, while %change_Glu/Glx_ was significant, with higher effect size from baseline compared to 3.5.1 (Hedge’s G_Glu/Glx-mean_ = 0.24, 95% CI: −0.066 – 0.553, I^2^ = 85.04, p = 0.045). GABA data show an overall lower effect for both mean and %change and did not reach statical significance, consistent with section 3.5.1.

Figure 8 shows data for Glu/Glx and GABA by stimulus domains across high-quality studies only. Several domains contained only a single high-quality study, therefore, results in domains such as stress (Glx and GABA) and motor (GABA) remained relatively the same. Mean_Glu_ shows a difference for the motor domain when only high-quality studies were included, indicating an increase of Glu-mean compared to the baseline condition (Hedge’s G_Glu-mean_ = 0.37, 95% CI: 0.004 – 0.743, I^2^ = 0). Exercise, learning, pain, and visual domain remained non-statistically significant for all metabolite types.

**Figure 8:**
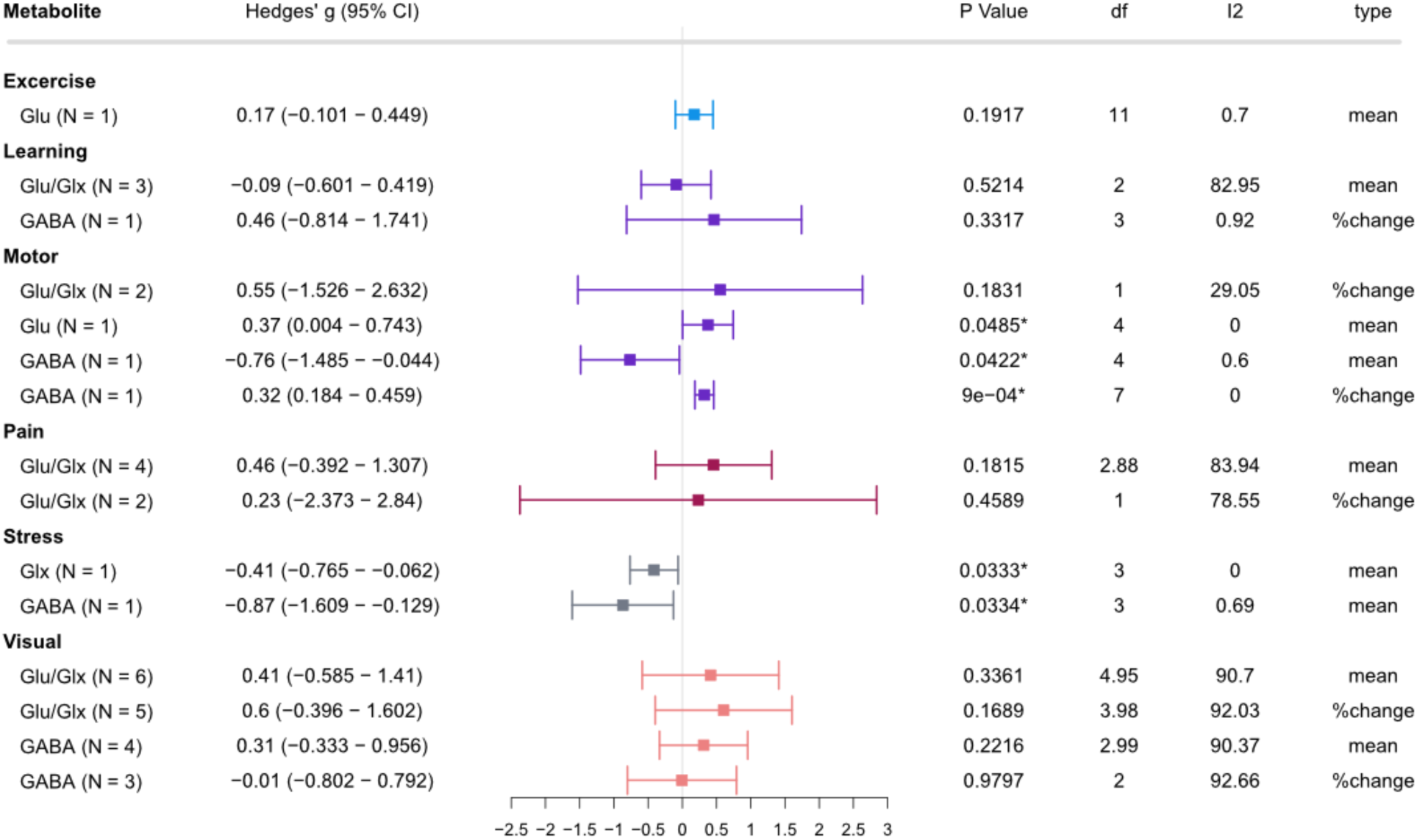
Meta-analysis of only ‘high quality’ studies as assessed by MRS-Q pooled based on stimulus domains. N: number of studies included; I^2^: I^2^ index for heterogeneity. A high I^2^ suggests there are external factors and biases driving the dispersions of effect sizes. *Statistically significant at p <0.05, and at p <0.01 when the degrees of freedom < 4 for RVE t-tests.

##### 3.6.2 Effect of number of transients and voxel size

First, we assessed whether effect size was correlated with the number of transients and voxel size. The number of transients mentioned here is the number of transients that was averaged across for metabolite quantification (e.g., per acquisition block or per one window width for sliding window analysis). There was statistically significant relationship between effect size and the number of transients for mean_Glu_ (ρ = −0.3, p = 0.0062). All other metabolites showed no significant relationship with number of transients (mean_GABA_: ρ = 0.021, p = 0.9, mean_Glx_: ρ = −0.27, p = 0.084). Percentage change in GABA (%change_GABA_: ρ = −0.21, p = 0.079), Glu (%change_Glu_: ρ = −0.15, p = 0.11), and Glx (%change_Glx_:ρ = −0.26, p = 0.2) showed no significant correlations between number of transients and effect size (Figure 9).

**Figure 9:**
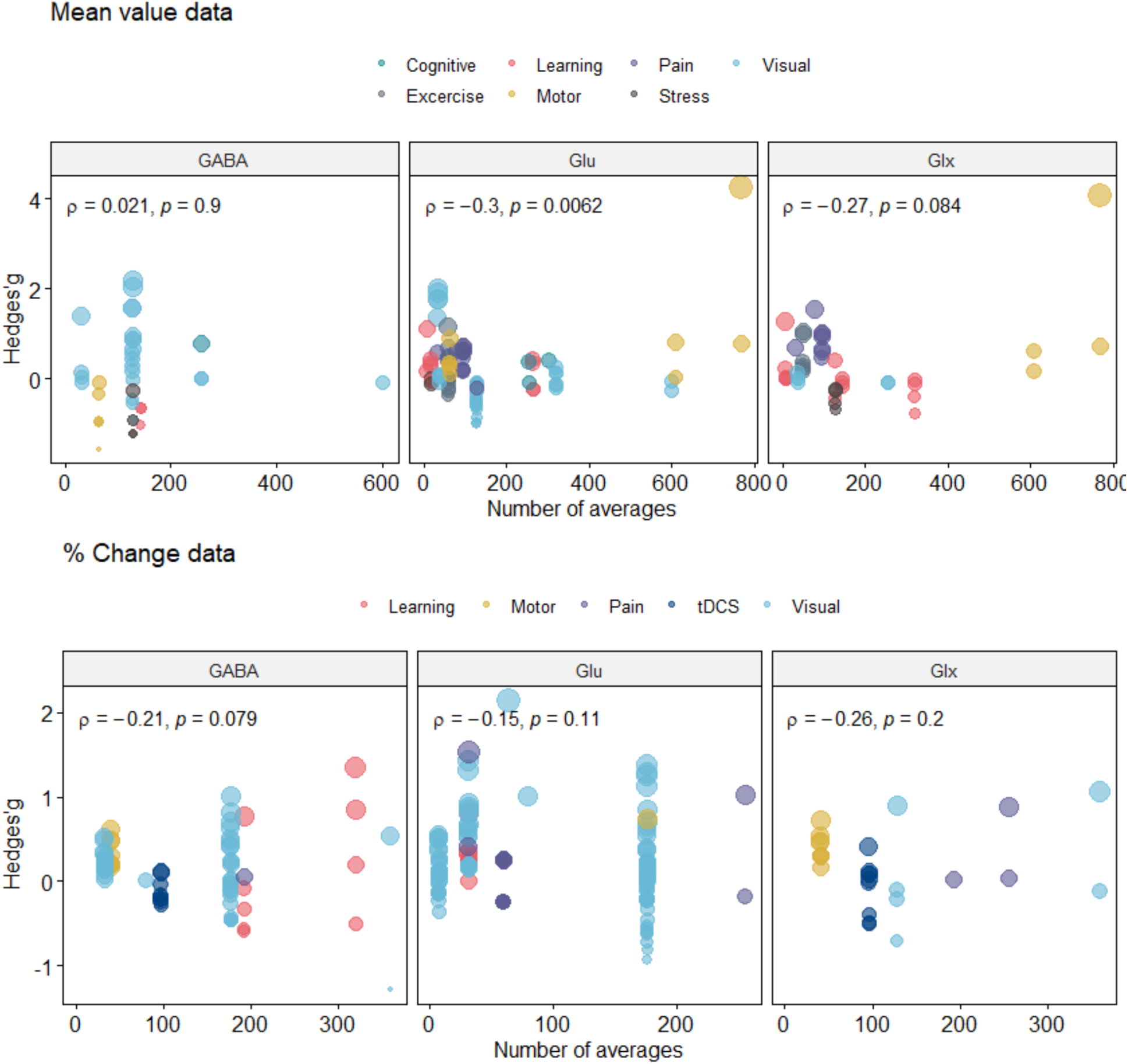
Relationship between effect size and number of transients used in included studies. The size of each dot represents the weight of effect size. met: metabolites; GABA: γ-Aminobutyric acid; Glu: Glutamate; Glx: Glutamine + Glutamate.

To analyse the association between effect size and number of transients, we binned studies based on the number of transients. Most of the studies used a number of transients in the range of 65-128 and 129-256 for metabolite quantification (n = 13 for each bin). A small increase in percentage GABA (Hedge’s G_GABA-%change_ = 0.23 – 0.32) and a small decrease in mean_GABA_ (Hedge’s G_GABA-mean_ = −0.76) were observed for studies with a limited number of transients (1-32 and 33-64). However, these significant results included data from only one study (Figure 10). The results were inconclusive when analysing these data by stimulus type, as only 1-4 studies were included for each stimulus type (Supplementary Figure 4).

**Figure 10:**
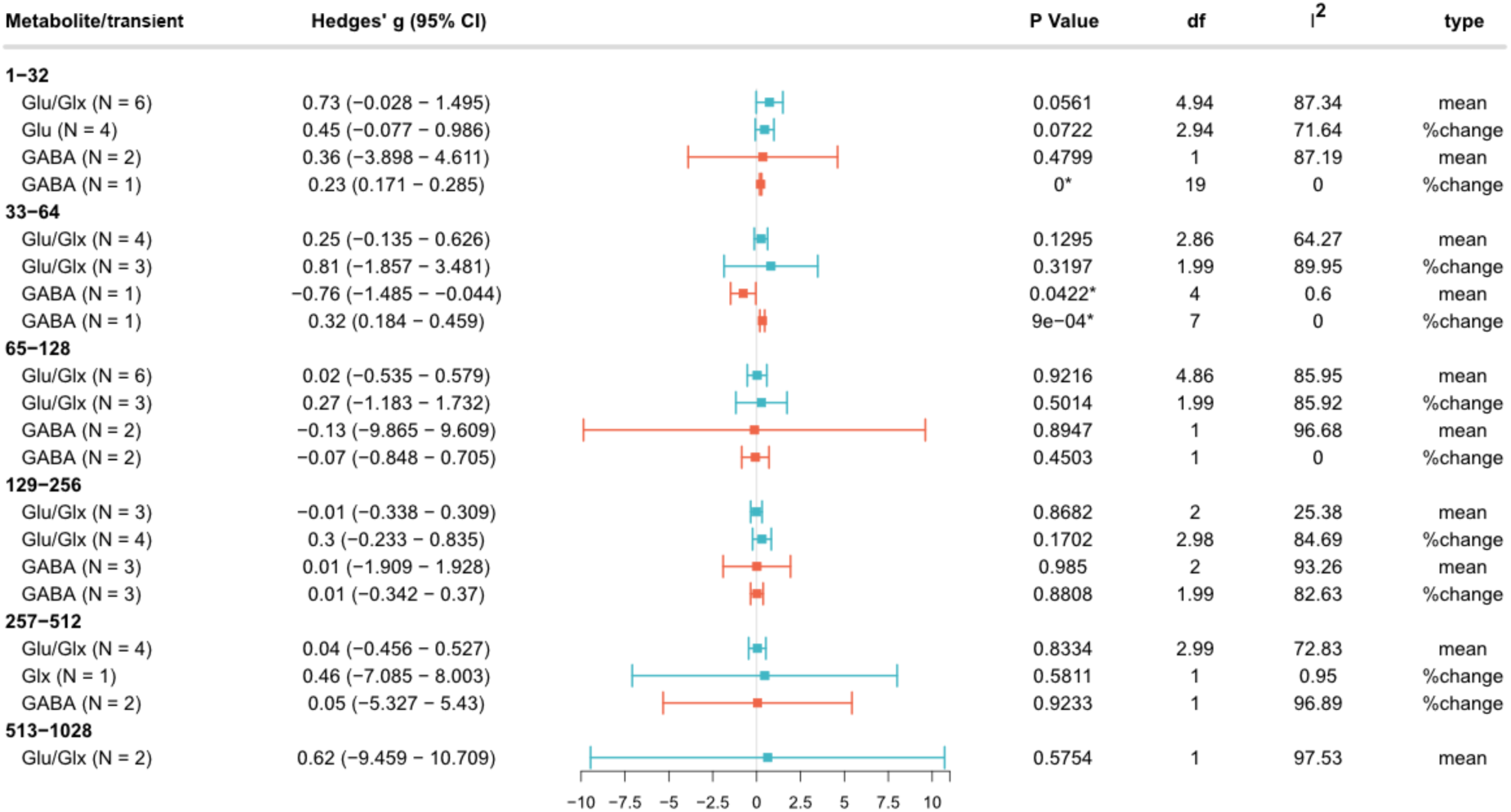
Influence of range of transient width on metabolite levels. N: number of studies included; I^2^: I^2^ index for heterogeneity. A high I^2^ suggests there are external factors and biases driving the dispersions of effect sizes. *Statistically significant at p <0.05, and at p <0.01 when the degrees of freedom < 4 for RVE t-tests.

The relationship between voxel size and effect size was different based on type of data (mean or %change) (Figure 11). For fMRS studies reported in mean metabolite levels, the effect sizes showed a negative relationship with voxel size (mean_GABA_: ρ = 0.42, p = 0.012; mean_Glu_: ρ = 0.066, p = 0.55; mean_Glx_: ρ = −0.42, p = 0.0059). Conversely, in studies reporting %change from the baseline condition, we observed a positive relationship between effect size and voxel size for all type of metabolite (%change_GABA_: ρ = 0.12, p = 0.3; (%change_Glu_: ρ = 0.19, p = 0.043; (%change_Glx_: ρ = 0.41, p = 0.039). Only mean_Glu_ and %change_GABA_ did not demonstrate a significant relationship with voxel size (Figure 11).

**Figure 11.**
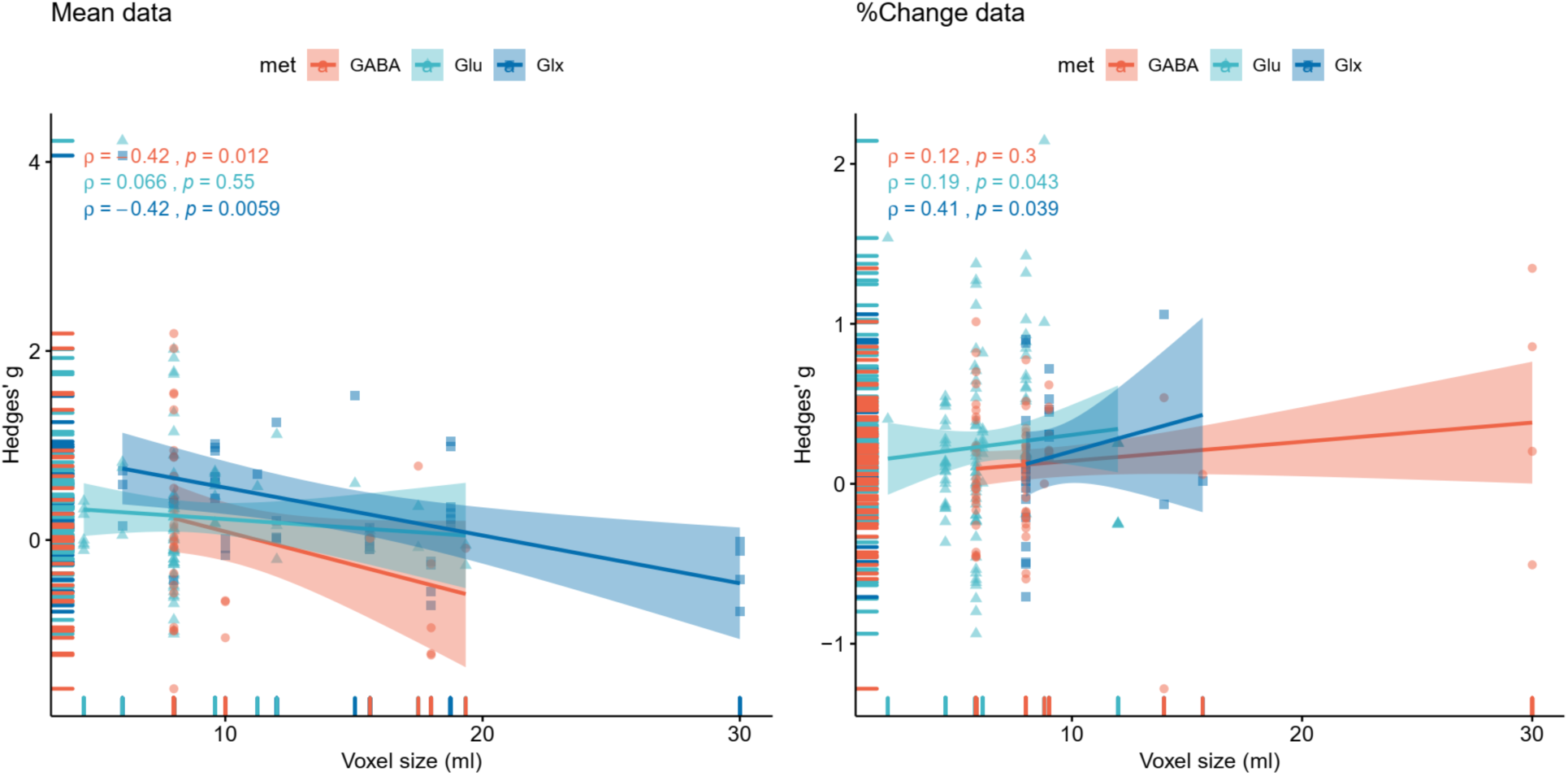
Effect size of each study in relation to voxel size in milliliters. The line represents a linear regression line for visual purposes only. ρ: Spearman’s rho; met: metabolites; GABA: γ-Aminobutyric acid; Glu: Glutamate; Glx: Glutamine+Glutamate.

## Discussion

### 1. Summary of the findings

We systematically evaluated and synthesized the fMRS literature on GABA and Glu/Glx to date (mid 2021). Overall, results show a wide variability in effect sizes and directionality for both Glu/Glx and GABA when generalized across design and stimulus domain. Most of the Glu/Glx studies showed positive trends (increases) during stimulation compared to baseline (at rest), while GABA studies generally showed negative trends (decreases) compared to baseline. The increase in Glu/Glx levels is in agreement with several animal studies showing an association between neuronal activation and Glu/Glx in response to task or stimuli (Just and Faber, 2019; Takado et al., 2021), which also correlates with BOLD signal activation (Just et al., 2013; Baslow et al., 2016; Just and Sonnay, 2017). Significant changes in Glu and Glx from baseline only had a small to average effect size (Hedge’s G_Glu and Glx_= 0.29 - 0.47). Although changes in GABA compared to baseline were not statistically significant across studies, the general directionality of *decreased* GABA levels is consistent with a previous narrative review by Duncan et al (2014) suggesting that GABA tends to be negatively correlated with task-evoked neuronal responses, as well as with studies showing that inhibition tends to decrease during repeated stimulation or learning (Stagg et al., 2011; Heba et al., 2016; Kolasinski et al., 2019). Ultimately, this meta-analysis shows that current fMRS works show large variety within domain and stimulus type, small effect sizes, and susceptibility to factors beyond experimenter control. While standardised reporting is becoming more widespread in MRS field, fMRS does not always adhere to the same principles and additional reporting standards need to be developed. This includes thorough reporting stimulus details and analysis methods, including open access to analysis code and stimulation paradigms, as these are likely driving the heterogeneity as well. This review revealed several important factors that need to be considered when performing and interpreting fMRS studies, which are detailed in the following sections.

### 2. Effect of fMRS design

#### 2.1 Effect of fMRS paradigm: block paradigm or event-related

In the current meta-analysis, the magnitude of effect sizes was observed to be smaller for block designs than event-related designs. This is in agreement with a previous meta-analysis of fMRS of Glu (Mullins, 2018). However, block designs provided more consistent results for Glu/Glx from tighter 95%CI of the averaged effect sizes compared to event-related designs, suggesting that block paradigms may be better at capturing Glu/Glx changes. On the other hand, event-related paradigms showed a wider range of confidence intervals compared to block design (event-related: 95% CI of −0.23 to 5.59, Block: 95% CI of −0.406 – 0.605). Although speculative, perhaps the most relevant difference between these two paradigms is that they are likely probing different brain processes, i.e., fast-acting neurochemical response through event-related designs and slower homeostatic processing or plasticity in block paradigms.

Block designs have the potential advantage of robust metabolite quantification as signal averaging is performed during a sustained stimulus. Habituation and adaptation to repeated stimulation with a potential summative effect likely plays a key role in block designs (Michels et al., 2012; Betina Ip et al., 2017; Ligneul et al., 2021). Signal averaging over a longer time course has been shown to smooth out any task-based dynamics of neural activity (Mangia et al., 2007; Mullins, 2018) and brain homeostasis during long stimulation blocks might lead to dismissal of, or minimal, metabolic changes (Mangia et al., 2012; Apšvalka et al., 2015).

These limitations can be overcome by time-locking fMRS to stimulus onset and assessing metabolic changes with higher temporal resolution. The temporal resolution of the event-related approach can be brought to under 30 seconds or less, allowing for measurement of a relatively fast response at the cost of increased measurement uncertainty of the individual time point due to decreased SNR. Several approaches have been implemented to successfully improve temporal resolution without sacrificing SNR, including sliding window, and/or averaging over participants, which will be discussed further in Section 3.1.

It is likely that the optimal choice of paradigm depends on the targeted stimulus domain. Any study with “long term” change (i.e., learning, memory, or even pharmacological approaches) may consider using block paradigms as these hold an advantage of higher SNR (Jahng et al., 2016; Bezalel et al., 2019; Vijayakumari et al., 2020). As previously discussed, block design often involves repeat stimulation with the theorised summation brain response, while event-related designs with fewer transients are likely to elicit a smaller response, which, even when averaged together, is not driven by repeated summation of stimuli. While this is not the right approach to assess transient responses, when someone is interested in more long-term changes, both our data and prior work suggests block designs may be more robust (Jahng et al., 2016; Bezalel et al., 2019; Vijayakumari et al., 2020). While this is speculative, our meta-analysis based on available data showed that block designs tend to have higher effect sizes than event-related designs. Nevertheless, careful fMRS paradigm design might allow for investigation of both block and event-related analysis within the same acquisition (Apšvalka et al., 2015; Stanley et al., 2017; Woodcock et al., 2018) through careful study design, but this is not widely used.

#### 2.2 Effect of stimulus domain

The directions and magnitudes of metabolic changes are influenced by stimulus domain. A significant increase compared to baseline was observed for Glu/Glx in five domains (exercise, learning, motor, stress, tDCs). Increased Glu/Glx during stimulation is in line with studies showing that neuronal responses require increased energy metabolism and/or excitatory neurotransmission. Although effect sizes were small, GABA concentrations tended to decrease in response to stimulation, except for %change in the motor domain. This is in agreement with previous studies demonstrating a negative relationship between regional neural activation and GABA, and a deactivation of GABAergic mechanisms when excitation is required (Duncan et al., 2014; Kiemes et al., 2021). It has been suggested that task-related GABA changes are more robustly observed in stimulus paradigms with a change in behavioural performance (Ip and Bridge, 2021), such as learning (Frangou et al., 2018, 2019), motor or sensory performance (Stagg et al., 2011; Heba et al., 2016; Kolasinski et al., 2019), and stress (Houtepen et al., 2017; Lynn et al., 2018a); this is reflected in our meta-analysis results, and GABA changes do not appear particularly robust. fMRS studies in pain appeared to be most consistent, but most domains show huge variation in their responses. GABA changes tend to be moderate at best and appear very domain- and approach, specific.

The high I^2^ across stimulus domains observed in this meta-analysis reflects the high degree of heterogeneity in results for different paradigms and stimuli even within stimulus domain. While we expected some variation as stimulus parameters and stimulation approach will differ between studies, we were surprised by this large heterogeneity. It should be noted that classification of stimulation domains may vary depending on individual opinion and judgement. For example, we grouped all visual stimulation fMRS studies into one category, despite differences in experimental design, stimulus intensity, and stimulus duration, which likely influenced the observed results (Mullins, 2018; Stanley and Raz, 2018; Ip and Bridge, 2021). Especially in the visual domain, we found a lot of heterogeneity, likely due to the variety in visual tasks including flashing checker boards with different flickering frequency, movie or clip-videos as visual stimulus, rotating checkerboard, and visual stimulations with variations in contrast level (Mangia et al., 2012; Kim et al., 2013b, 2014; Betina Ip et al., 2017; Mekle et al., 2017; Bednařík et al., 2018; Martínez-Maestro et al., 2019). Previous studies demonstrated regional cerebral blood flow change in linear function with stimulus repetition rates that peaked at approximately 8 Hz then decline above this frequency (Fox and Raichle, 1984; Bejm et al., 2019). Previous fMRI studies also reported BOLD response to be depends on stimulus patterns (Krüger et al., 1998; Hoge et al., 1999). Similarly, perhaps approaches with higher SNR (such as 7T) are more sensitive to changes (Mangia et al., 2012).

Combining visual stimulus studies was necessary, however, as separating them out further would lead to single study analysis, which is not particularly useful for meta-analytical purposes. However, we do know stimulus parameters can have different effects. Previous studies have demonstrated a lack of both Glu and BOLD signal changes at low visual contrast level, whereas only high stimulus intensity elicited a measurable and significant Glu response (Ip et al., 2019). This suggests that stimuli used for fMRS are preferably ones with high-intensity to evoke a sufficiently salient response (e.g., in a considerable number of neurons) to cause neurometabolite production or spillover (Yashiro et al., 2005; Gonçalves-Ribeiro et al., 2019), which leads to a measurable transient change that can be measured with MRS. Additionally, MRS-derived neurometabolite signals are non-specific and reflect all cellular component (e.g., cytosol, extracellular space, vesicle, synaptic cleft, etc.). It is possible that a smaller brain response with less SNR (e.g., one induced by repetitive stimulation) could be masked by other metabolic responses with higher SNR (e.g., energy usage, steady state).

#### 2.3 Effect of ROI

Although we intended to study the effect of ROI on effect size, there was insufficient data to draw firm conclusions. Despite the occipital ROI being the most studied ROI in fMRS (and MRS in general, (Puts and Edden, 2012)), and with the benefit of high-quality spectra due to its homogenous field relative to other ROIs (Juchem and de Graaf, 2017), only %change_Glu/Glx_ was significantly increased compared to baseline. A significant *increase* of GABA was demonstrated for frontal and parietal ROIs which included fMRS studies of visual, exercise, motor, stress and learning stimulus; all these involved some kind of repeated stimulation and likely to reflect plasticity. This is consistent with the notion that both frontal and parietal regions play important roles in regulating inhibitory control of behaviour (Aron et al., 2004; Narayanan and Laubach, 2017; Hermans et al., 2018). An increase in Glu/Glx was demonstrated for insular cortex and other temporal lobe regions. While we can only speculate why this appears to be more robust t, it might be that there is less variation in the approach used for insular regions compared to other regions, for example, visual studies. It is possible that paradigms targeting insular/parietal regions elicit stronger responses in these regions than visual stimuli do in visual regions, but it might also be the case that voxels have less heterogeneity (as heterogeneity even within occipital lobe is large, and different occipital regions have very different roles).

The differences in both of direction and magnitude due to anatomical differences and functional differences of ROIs are not surprising (Gordon et al., 2017; Zhang et al., 2020). Different brain regions typically contain different tissue compositions (i.e., white and grey matter) (Pouwels and Frahm, 1998; Amaral et al., 2013). Differences in tissue composition also leads to variation in metabolism with grey matter having higher energy consumption compared to white matter (Amaral et al., 2013; Ford and Crewther, 2016), which in turn, affects GABA, Glu and Glx levels (Rae et al., 2009; Rae, 2014) *and see also next section*. We were not able to determine the role of tissue composition and subsequent partial volume correction, which accounts for much variation in the estimation of GABA and Glx/Glu, due to limited available and reported data. Another possible explanation for differences in effect sizes between ROIs could arises from increase SNR in certain regions (e.g., occipital lobe) with close proximity to the receiver coil as well (Di Costanzo et al., 2007; Minati et al., 2010). Nevertheless, further primary studies are required to further elucidate the relationship between effect sizes and brain region.

#### 2.4 Possible mechanisms underly metabolite changes

While directional changes in the neurometabolite responses were observed in this meta-analysis, the mechanisms underlying these changes remain unclear. Metabolite concentrations obtained in fMRS studies originate from all cell compartments (i.e., cell body, cytosol, synaptic cleft, etc.) (Puts and Edden, 2012). The brain’s response to external stimuli consists of a complex interplay between neuronal mechanisms. This includes changes in blood flow, changes in neurotransmitter transport, production and breakdown, and brain oxidative metabolism (Fox and Raichle, 2007; Mangia et al., 2009; Takado et al., 2021). Besides neuronal synaptic activity, metabolic processes also contribute to the neurometabolite levels measured in MRS (e.g., the TCA cycle)(Dienel, 2012; Magistretti and Allaman, 2015).

Our finding of increased Glu/Glx during stimulation/tasks is in agreement with several studies that link Glu and brain responses to stimulus such as perception, visual activation, motor activation, learning, and memory (Gao et al., 2013; Magalhães et al., 2019; Ligneul et al., 2021). Glu plays a major role during activity-dependent energy demands as the most abundant amino-acid and the main excitatory neurotransmitter in the brain (Ligneul et al., 2021). Increasing evidence demonstrates the close regulation between glucose consumption and glutamate-glutamine cycling (Sibson et al., 1998; Rothman et al., 2003), which was theorised to lead to increasing Glu levels during the BOLD-activation period (Betina Ip et al., 2017; Vijayakumari et al., 2018; Martínez-Maestro et al., 2019). Additionally, Glu is also a major determinant for neuronal plasticity during periods of high neural activity as Glu influence the production of of brain-derived neurotrophic factor (BDNF) which regulates survival, differentiation and synaptogenesis in the CNS to change patterns of neuronal connectivity (Gonçalves-Ribeiro et al., 2019; Valtcheva and Venance, 2019). Indeed, Glu release by neurons and its uptake to astrocytes for recycling via glutamine is thought to represent 70-80% of total brain glucose consumption (Hertz and Rothman, 2016). That said, it is not possible to differentiate metabolic Glu from vesicular or synaptic Glu, and caution in the interpretation of Glu/Glx changes is important; one cannot simply extrapolate these changes to changes in neurotransmission.

Previous studies have demonstrated the relationship between GABA as measured with MRS and the gene encoding for glutamic acid decarboxylase (GAD) 67. GAD 67 is responsible for converting Glu into GABA under baseline conditions and the majority of GABA production, and is present in both cell bodies (Marenco et al., 2010) and synapses. Therefore, MRS quantified GABA is often said to reflect ‘inhibitory tone’ (Rae, 2014; Peek et al., 2020). The relationship between GABA and neuronal activation (or deactivation) is less consistent, and often dependent on the task used. Previous work has shown that increased GABA levels are associated with increased BOLD signal in response to an interference task (Kühn et al., 2016) and in response to pharmacological manipulation in rat brain (Chen et al., 2005). However, other studies have shown that higher baseline GABA was associated with lower BOLD response amplitude (Muthukumaraswamy et al., 2012; Rae, 2014; Stanley and Raz, 2018). It has been suggested that Glu/Glx and GABA changes in response to stimulation comprise of both energy usage and neural process facilitating a shift into new metabolic steady-state *by* shifting the excitation/inhibition equilibrium, linking these two processes more directly (Just et al., 2013; Lynn et al., 2018b). A recent fMRS study in animals models showed that increases in GABA after repeated tactile stimulation were consistent with two-photon microscopy measures of increased inhibitory activity, and increases in Glu with increased excitatory activity, suggesting that functional changes in GABA and Glu measured through MRS are indeed reflective of increased inhibitory neurotransmission (Takado et al., 2021).

### 3. Effect of fMRS parameters

Beyond assessing fMRS through differences and changes in the ‘bulk’ metabolite response to stimulation, it is also important to investigate differences at the level of acquisition and analysis. In this meta-analysis, we demonstrated the effect of MRS parameters such as number of transients, voxel size, timing, and MRS quality limitations on reported metabolite concentrations.

#### 3.1 Number of transients

The results reported in our meta-analysis illustrate the variability of methods used in fMRS studies. For both Glu/Glx and GABA, effect sizes seem to be higher for a lower number of transients, and effect size decreases as the number of transients increase. There are several possible explanations for this; One is that low transient sizes lead to lower SNR and unreliable spectral quantification (Mikkelsen et al., 2018), which potentially lead to biased metabolite concentration changes. Another possible explanation is that rapid changes in the first few minutes due to neurotransmitter release might influence the effect sizes observed with a small number of transients due to their higher temporal resolution (Mullins, 2018; Ligneul et al., 2021). On the other hand, a larger number of transients might lead to lower effect size observed due the effect being averaged out over a longer period of time (Ip and Bridge, 2021), thus diluting any rapid changes. Ultimately, conclusions are difficult to draw without a measurable ground truth, since the spectral fitting process itself may introduce quantitative bias depending on SNR (and therefore the width of the averaging window). Synthetic simulated data can be useful to elucidate the accuracy, precision, and biases of spectral fitting when attempting to resolve small temporal changes.

Given the approximate 10^4^ times lower metabolite concentrations relative to water, and thus low SNR, spectra are often collected with long acquisition times. These acquisition times are often longer than the assumed temporal dynamics with fast metabolite changes in less than 1s (Apšvalka et al., 2015; Bednařík et al., 2015; Mullins, 2018; Ligneul et al., 2021) which likely reflect changes in visibility in existing metabolite pools. Several spectral averaging methods have been applied to overcome this trade-off between temporal resolution and SNR (Kanowski et al., 2004; Mikkelsen et al., 2018). One of these averaging methods included averages fMRS data across short sequential acquisition blocks (Kolasinski et al., 2019). Others used time-locking to stimulus onset followed by averaged transients acquired during stimulus presentation or baseline, comparing the two, as event-related averaging (Lally et al., 2014 p.201; Apšvalka et al., 2015; Stanley et al., 2017). Some studies have averaged across a small number of transients but across participants to obtain group-level spectra with higher temporal resolution (Apšvalka et al., 2015; Bednařík et al., 2015; Fernandes et al., 2020). Others have applied a ‘sliding window’ or ‘moving averages’ approach (i.e., average transients in blocks then shifting the averaging over time by a certain transient window width) to detect a dynamic trace of metabolite changes (Mangia et al., 2007; Schaller et al., 2013; Fernandes et al., 2020; Rideaux, 2020).

Our results are in agreement with studies suggesting averaging across a small number of transients has an advantage of higher temporal resolution for detecting rapid modulation of metabolite levels (Lally et al., 2014; Betina Ip et al., 2017; Ligneul et al., 2021). A longer averaging window might be better associated with moving towards a new steady metabolism as described above (Betina Ip et al., 2017; Lynn et al., 2018a). Furthermore, the brain likely responds differently to different types of stimuli, and in a region-specific manner, once again emphasising that task design needs to be tailored towards the question of interest. Surprisingly, we know very little about the actual temporal dynamics of these metabolites thus makes it difficult to a priori choose the best acquisition strategy. Only a few studies were included to allow for the consideration of the impact of transient width, which supports the urgent needs in of more primary studies of fMRS with varying time windows.

#### 3.2 fMRS timing

Our analysis allowed us to explore whether effect sizes change with time of acquisition. While exploratory, these time-resolved fluctuation patterns suggest different response functions for different brain regions or stimulus domains, *and* between GABA and Glu/Glx. Some studies observed a fast Glu response early in a working memory task, but not later in the task (Woodcock et al., 2018), while others observed Glu reaching a new steady state 1 to 2 minutes after stimulus onset (Mangia et al., 2007; Schaller et al., 2013). Previous studies of GABA and Glx in response to visual stimulation demonstrated concentration drifts over time in opposite directions while participants were at ‘rest’ before stabilising (in steady state) after around 500 seconds (Rideaux, 2020). As discussed in previous sections, the time courses of neurometabolites in response to stimulus domain are a topic of great interest and require further elucidation. This perhaps can be achieved by varying the time of fMRS acquisition and stimulus onset in high-field MR at > 3 T, while aiming for the best temporal resolution possible.

#### 3.3 Others MR-instrument-related limitations

fMRS is also sensitive to other instrument and acquisition-related limitations. MRS offers low spatial specificity as large voxels (often >15 ml) are required for sufficient SNR. Reducing voxel size requires increasing acquisition time to maintain SNR, which is not only impractical, but also increases the risk of scanner drift and participant motion, especially in clinically sensitive motion-prone groups such as prenatal and people with neurodevelopmental conditions (Mikkelsen et al., 2018; Hui et al., 2021; Ip and Bridge, 2021).

### 4. Quality assurance of MRS

Differences in fMRS parameters go hand-in-hand with quality assurance. There is no consensus on minimally best practice for fMRS to date. Currently available quality assurance and reporting metrics (MRS-Q and MRSinMRS) were designed for static MRS (Peek et al., 2020; Lin et al., 2021), and do not take into account functional approaches where the averaged number of transients is often lower to achieve better temporal resolution. Notably, many studies reported here used smaller voxel sizes compared to consensus recommendation (~27 ml for edited MRS for GABA, 3 T, and ~3.4 ml for unedited at 128 transients, 3T) (Lin et al., 2021). Smaller voxel size inconsistent with consensus standards was often observed in particular for spectral editing of GABA, and findings may be less reliable due to insufficient SNR. Here, we used standard language for quality assessment (such as high or low quality) but should of course note that this language often refers to studies not reporting sufficient information. It is our hope that with the increasing consensus in reporting, this will become less of a concern. We should also note that some studies used “low quality” approaches compared to the consensus now but need to see these in a historical perspective. Despite several studies reporting inadequate fMRS parameters, our sensitivity analysis based on study quality shows no extreme changes from analyses including all studies. While there is room for improvement for reporting of fMRS, most of the studies used adequate fMRS scan parameters. It is possible that the number of transients is less important when modelling time-course data and using within-participant designs. Establishing minimum reporting standard in this early stage would greatly increase reproducibility in a field that offers an almost unlimited number of data analysis strategies.

### 5. Sources of bias

As discussed in the previous sections, there are various sources of bias in fMRS study design, acquisition, and analysis parameters (i.e., brain area, voxel size, number of transients, and metabolite unit, e.g., percentage change or mean concentration). Study quality assessments further suggest that fMRS studies lack randomisation and blinding of participants. Additional risk of bias could arise from selection of participants, for example, studies often using colleagues as participants for the study. fMRS studies such as stress or pharmacological designs often use a pre-post within-participant design, this introduces bias into the analysis (Ma et al., 2020) and potentially leads to reporting of positive results (publication bias) (Rosenthal, 1979; Murphy and Aguinis, 2019). Additional sources of bias were beyond the scope of this meta-analysis. These include the general experiment design, such as population sampling and type of baseline condition such as difference type of visual baseline condition of eyes close or a fixation cross (Ip et al., 2019; Ip and Bridge, 2021), the choice of analysis approach including differences between spectral modelling algorithms (Zöllner et al., 2021, 2022; Craven et al., 2022; Marjańska et al., 2022), quantification and referencing (metabolite in institutional units [IU], absolute concentration or ratio to creatine [Cr], etc.)(Porges et al., 2017), and how results are reported (e.g., reported only in percentage change but not in concentration). In particular, the choice of quantification reference compound might have a strong impact, although it is assumed typical reference compounds such as Cr or NAA are unlikely to change with stimulation (Wilson et al., 2019). One important parameter that needs further investigation and consensus treatment is linewidth adjustment based on the BOLD signal. Since haemoglobin is paramagnetic when deoxygenated, but diamagnetic when oxygenated, local magnetic susceptibility depends on the blood-oxygen level. BOLD activation causes narrowed MRS lines and increased signal magnitude that can lead to overestimation of metabolite levels if uncorrected (Zhu and Chen, 2001; Betina Ip et al., 2017). In the present study, we did not have sufficient information to perform an analysis on these topics.

As shown in the results, the type of data reported (mean or %change) influences the effect sizes observed. Most fMRS studies reported results as %change, followed by ratio to reference molecule (e.g., tCr, NAA). Our meta-analysis avoids the secondary calculation of data by analysing data as presented. For the sake of transparency and understanding these impacts, we suggest reporting data in both comparative result (e.g., %change, change from baseline) and in mean metabolite concentration (e.g., ratio to reference metabolite, mmol/kg, institutional units) in the future. These results often support and strengthen each other and increase comparability between studies. Future study could investigate the Glu and GABA ratio as a theoretical index of E/I balanced, although the exact relationship between MRS-derived Glu/GABA concentration and E/I balanced is still under active debate (Steel et al., 2020; Rideaux, 2021).

### 6. Limitation of the current meta-analysis

Several limitations to this meta-analysis study need to be acknowledged when interpreting this current work. While the data were considered based on data type (mean and %change), we had to assume that the units included were on the same scale, and that reference metabolites concentration (e.g., creatine or NAA) remained relatively unchanged (Steen et al., 2005; Rae, 2014). Another potential limitation is that we included all studies regardless of study quality as assessed by both ROBANS and MRS-Q in our main meta-analysis. The sensitivity analysis of high-quality studies according to MRS-Q suggested that the studies included showed interchangeable results regardless of quality, and we recognize that the MRS-Q, while useful, does not fully apply to functional MRS. While we aim to comprehensively include all fMRS studies to date, unfortunately our search strategy may have missed out on more recent work, such as Ip et al. (2019); we did not identify this paper through other means. Lastly, there was a lack of statistical power for some stimulus domains and fMRS paradigm due to the small numbers of studies included. Taken together, fMRS is a field with enormous possibility, but with several sources of bias and variability that need to be addressed.

In this current study, we employed the RVE method to synthesize effect sizes for each stimulus domain from the multiple outcomes available (i.e., multiple within-study outcomes). While some significant changes from baseline were noted in some stimulus domains, often they led to single-study meta-analyses with several datasets from various timepoints included (varying from 3-9) in each single-study per each domain. These results therefore need to be interpreted with care as there is study-bias.

### 7. Conclusion

We established effect sizes and directionality of the GABA, Glx and Glu response in all currently available fMRS studies. Our results demonstrated relatively small effect sizes and large heterogeneity, limiting the current state of fMRS as a technique in investigating neurodynamic responses in the healthy brain. However, we attempt to address these limitations and hope that advances in these approaches have promise for application in atypical brain function. fMRS of clinical conditions is surprisingly under-studied, but holds promise for understanding a dynamic system, with potential implications for drug response and diagnosis. As such, fMRS holds great potential to be used alongside other techniques to perturb GABA and Glutamate mechanisms, including TMS and pharmacological challenges and assess the impact on the system in both typical and atypical brain. Furthermore, combining fMRS with other imaging techniques, such as EEG or fMRI, allows for associating (f)MRS with distinct neural mechanisms associated with E/I balance.

This meta-analysis highlights the urgent need for consensus for standardised reporting and minimal best practices to improve the reproducibility of fMRS. Additionally, there remains a lack of fundamental knowledge of fMRS, for example, with respect to metabolic time courses. Establishing fMRS paradigms and parameters that evoke metabolic responses with high reliability and reproducibility would be of great interest in this early state of the field as it would allow for measuring atypical responses more readily, and ultimately lead to elucidation of underlying mechanisms of brain function in both health and disease.

## Acknowledgments

DP is funded by a Chang Phueak Scholarship, Chiang Mai University, ChiangMai, Thailand. GO receives salary support from R00 AG062230 and R21 EB033516.

## References

Amaral A, Meisingset T, Kotter M, Sonnewald U (2013) Metabolic Aspects of Neuron-Oligodendrocyte-Astrocyte Interactions. Frontiers in Endocrinology 4 Available at: https://www.frontiersin.org/article/10.3389/fendo.2013.00054 [Accessed January 25, 2022].

Ankit Rohatgi (2021) WebPlotDigitizer. Available at: https://automeris.io/WebPlotDigitizer.

Apšvalka D, Gadie A, Clemence M, Mullins PG (2015) Event-related dynamics of glutamate and BOLD effects measured using functional magnetic resonance spectroscopy (fMRS) at 3T in a repetition suppression paradigm. NeuroImage 118:292–300.

Archibald J, MacMillan EL, Graf C, Kozlowski P, Laule C, Kramer JLK (2020) Metabolite activity in the anterior cingulate cortex during a painful stimulus using functional MRS. Scientific reports 10:19218–19218.

Aron AR, Robbins TW, Poldrack RA (2004) Inhibition and the right inferior frontal cortex. Trends in Cognitive Sciences 8:170–177.

Baeshen A, Wyss PO, Henning A, O’Gorman RL, Piccirelli M, Kollias S, Michels L (2020) Test–Retest Reliability of the Brain Metabolites GABA and Glx With JPRESS, PRESS, and MEGA-PRESS MRS Sequences in vivo at 3T. Journal of Magnetic Resonance Imaging 51:1181–1191.

Baslow MH, Cain CK, Sears R, Wilson DA, Bachman A, Gerum S, Guilfoyle DN (2016) Stimulation-induced transient changes in neuronal activity, blood flow and N-acetylaspartate content in rat prefrontal cortex: a chemogenetic fMRS-BOLD study. NMR in Biomedicine 29:1678–1687.

Bednařík P, Tkáč I, Giove F, DiNuzzo M, Deelchand DK, Emir UE, Eberly LE, Mangia S (2015) Neurochemical and BOLD responses during neuronal activation measured in the human visual cortex at 7 Tesla. Journal of cerebral blood flow and metabolism : official journal of the International Society of Cerebral Blood Flow and Metabolism 35:601–610.

Bednařík P, Tkáč I, Giove F, Eberly LE, Deelchand DK, Barreto FR, Mangia S (2018) Neurochemical responses to chromatic and achromatic stimuli in the human visual cortex. J Cereb Blood Flow Metab 38:347–359.

Begg CB, Mazumdar M (1994) Operating Characteristics of a Rank Correlation Test for Publication Bias. Biometrics 50:1088–1101.

Bejm K, Wojtkiewicz S, Sawosz P, Perdziak M, Pastuszak Z, Sudakou A, Guchek P, Liebert A (2019) Influence of contrast-reversing frequency on the amplitude and spatial distribution of visual cortex hemodynamic responses. Biomed Opt Express 10:6296–6312.

Betina Ip I, Berrington A, Hess AT, Parker AJ, Emir UE, Bridge H (2017) Combined fMRI-MRS acquires simultaneous glutamate and BOLD-fMRI signals in the human brain. Neuroimage 155:113–119.

Bezalel V, Paz R, Tal A (2019) Inhibitory and excitatory mechanisms in the human cingulate-cortex support reinforcement learning: A functional Proton Magnetic Resonance Spectroscopy study. NeuroImage 184:25–35.

Boillat Y, Xin L, van der Zwaag W, Gruetter R (2020) Metabolite concentration changes associated with positive and negative BOLD responses in the human visual cortex: A functional MRS study at 7 Tesla. Journal of cerebral blood flow and metabolism : official journal of the International Society of Cerebral Blood Flow and Metabolism 40:488–500.

Borenstein M, Hedges LV, Higgins JP, Rothstein HR (2021) Introduction to meta-analysis. John Wiley & Sons.

Bottomley PA (1984) Selective volume method for performing localized NMR spectroscopy. Google Patents.

Bottomley PA (1987) Spatial localization in NMR spectroscopy in vivo. Annals of the New York Academy of Sciences 508:333–348.

Bowden J, Davey Smith G, Burgess S (2015) Mendelian randomization with invalid instruments: effect estimation and bias detection through Egger regression. International journal of epidemiology 44:512–525.

Cai K, Nanga RP, Lamprou L, Schinstine C, Elliott M, Hariharan H, Reddy R, Epperson CN (2012) The Impact of Gabapentin Administration on Brain GABA and Glutamate Concentrations: A 7T 1H-MRS Study. Neuropsychopharmacol 37:2764–2771.

Chen C, Sigurdsson HP, Pépés SE, Auer DP, Morris PG, Morgan PS, Gowland PA, Jackson SR (2017) Activation induced changes in GABA: Functional MRS at 7T with MEGA-sLASER. NeuroImage 156:207–213.

Chen W, Novotny EJ, Zhu XH, Rothman DL, Shulman RG (1993) Localized 1H NMR measurement of glucose consumption in the human brain during visual stimulation. Proc Natl Acad Sci U S A 90:9896–9900.

Chen Z, Silva AC, Yang J, Shen J (2005) Elevated endogenous GABA level correlates with decreased fMRI signals in the rat brain during acute inhibition of GABA transaminase. J Neurosci Res 79:383–391.

Chiappelli J, Shi Q, Wijtenburg SA, Quiton R, Wisner K, Gaston F, Kodi P, Gaudiot C, Kochunov P, Rowland LM, Hong LE (2018) Glutamatergic Response to Heat Pain Stress in Schizophrenia. Schizophr Bull 44:886–895.

Cleve M, Gussew A, Reichenbach JR (2015) In vivo detection of acute pain-induced changes of GABA+ and Glx in the human brain by using functional 1H MEGA-PRESS MR spectroscopy. NeuroImage 105:67–75.

Cleve M, Gussew A, Wagner G, Bär K-J, Reichenbach JR (2017) Assessment of intra- and inter-regional interrelations between GABA+, Glx and BOLD during pain perception in the human brain - A combined (1)H fMRS and fMRI study. Neuroscience 365:125–136.

Coghlan S, Horder J, Inkster B, Mendez MA, Murphy DG, Nutt DJ (2012) GABA system dysfunction in autism and related disorders: from synapse to symptoms. Neurosci Biobehav Rev 36:2044–2055.

Coxon JP, Cash RFH, Hendrikse JJ, Rogasch NC, Stavrinos E, Suo C, Yücel M (2018) GABA concentration in sensorimotor cortex following high-intensity exercise and relationship to lactate levels. The Journal of Physiology 596:691–702.

Craven AR, Bhattacharyya PK, Clarke WT, Dydak U, Edden RAE, Ersland L, Mandal PK, Mikkelsen M, Murdoch JB, Near J, Rideaux R, Shukla D, Wang M, Wilson M, Zöllner HJ, Hugdahl K, Oeltzschner G (2022) Comparison of seven modelling algorithms for γ-aminobutyric acid–edited proton magnetic resonance spectroscopy. NMR in Biomedicine 35 Available at: https://onlinelibrary.wiley.com/doi/10.1002/nbm.4702 [Accessed August 26, 2022].

Deelchand DK, Marjańska M, Henry P-G, Terpstra M (2021) MEGA-PRESS of GABA+: Influences of acquisition parameters. NMR in Biomedicine 34:e4199.

Dennis A, Thomas AG, Rawlings NB, Near J, Nichols TE, Clare S, Johansen-Berg H, Stagg CJ (2015) An Ultra-High Field Magnetic Resonance Spectroscopy Study of Post Exercise Lactate, Glutamate and Glutamine Change in the Human Brain. Front Physiol 6 Available at: https://www.frontiersin.org/articles/10.3389/fphys.2015.00351/full [Accessed June 20, 2021].

Di Costanzo A, Trojsi F, Tosetti M, Schirmer T, Lechner SM, Popolizio T, Scarabino T (2007) Proton MR spectroscopy of the brain at 3 T: an update. Eur Radiol 17:1651–1662.

Dienel GA (2012) Brain Lactate Metabolism: The Discoveries and the Controversies. J Cereb Blood Flow Metab 32:1107–1138.

Donahue MJ, Near J, Blicher JU, Jezzard P (2010) Baseline GABA concentration and fMRI response. Neuroimage 53:392–398.

Duarte JMN, Lei H, Mlynárik V, Gruetter R (2012) The neurochemical profile quantified by in vivo 1H NMR spectroscopy. NeuroImage 61:342–362.

Duncan NW, Wiebking C, Northoff G (2014) Associations of regional GABA and glutamate with intrinsic and extrinsic neural activity in humans—A review of multimodal imaging studies. Neuroscience & Biobehavioral Reviews 47:36–52.

Duval S, Tweedie R (2000) A nonparametric “trim and fill” method of accounting for publication bias in meta-analysis. Journal of the american statistical association 95:89–98.

Dwyer GE, Craven AR, Bereśniewicz J, Kazimierczak K, Ersland L, Hugdahl K, Grüner R (2021) Simultaneous Measurement of the BOLD Effect and Metabolic Changes in Response to Visual Stimulation Using the MEGA-PRESS Sequence at 3 T. Frontiers in Human Neuroscience 15 Available at: https://www.frontiersin.org/article/10.3389/fnhum.2021.644079.

Edden RAE, Barker PB (2007) Spatial effects in the detection of gamma-aminobutyric acid: improved sensitivity at high fields using inner volume saturation. Magn Reson Med 58:1276–1282.

Edden RAE, Muthukumaraswamy SD, Freeman TCA, Singh KD (2009) Orientation Discrimination Performance Is Predicted by GABA Concentration and Gamma Oscillation Frequency in Human Primary Visual Cortex. J Neurosci 29:15721–15726.

Egger M, Smith GD, Schneider M, Minder C (1997) Bias in meta-analysis detected by a simple, graphical test. BMJ 315:629–634.

Faghihi R, Zeinali-Rafsanjani B, Mosleh-Shirazi M-A, Saeedi-Moghadam M, Lotfi M, Jalli R, Iravani V (2017) Magnetic Resonance Spectroscopy and its Clinical Applications: A Review. J Med Imaging Radiat Sci 48:233–253.

Ferguson BR, Gao W-J (2018) PV Interneurons: Critical Regulators of E/I Balance for Prefrontal Cortex-Dependent Behavior and Psychiatric Disorders. Frontiers in Neural Circuits 12 Available at: https://www.frontiersin.org/articles/10.3389/fncir.2018.00037 [Accessed October 8, 2022].

Fernandes CC, Lanz B, Chen C, Morris PG (2020) Measurement of brain lactate during visual stimulation using a long TE semi-LASER sequence at 7 T. NMR in biomedicine 33:e4223–e4223.

Floyer-Lea A, Wylezinska M, Kincses T, Matthews PM (2006) Rapid Modulation of GABA Concentration in Human Sensorimotor Cortex During Motor Learning. Journal of Neurophysiology 95:1639–1644.

Ford TC, Crewther DP (2016) A Comprehensive Review of the 1H-MRS Metabolite Spectrum in Autism Spectrum Disorder. Frontiers in Molecular Neuroscience 9 Available at: https://www.frontiersin.org/article/10.3389/fnmol.2016.00014 [Accessed January 25, 2022].

Fox MD, Raichle ME (2007) Spontaneous fluctuations in brain activity observed with functional magnetic resonance imaging. Nat Rev Neurosci 8:700–711.

Fox PT, Raichle ME (1984) Stimulus rate dependence of regional cerebral blood flow in human striate cortex, demonstrated by positron emission tomography. Journal of Neurophysiology 51:1109–1120.

Frahm J al, Bruhn H, Gyngell M, Merboldt K, Hänicke W, Sauter R (1989) Localized high-resolution proton NMR spectroscopy using stimulated echoes: initial applications to human brain in vivo. Magnetic resonance in medicine 9:79–93.

Frangou P, Correia M, Kourtzi Z (2018) GABA, not BOLD, reveals dissociable learning-dependent plasticity mechanisms in the human brain King AJ, Silver M, Dosher BA, Greenhouse I, Silver M, eds. eLife 7:e35854.

Frangou P, Emir UE, Karlaftis VM, Nettekoven C, Hinson EL, Larcombe S, Bridge H, Stagg CJ, Kourtzi Z (2019) Learning to optimize perceptual decisions through suppressive interactions in the human brain. Nat Commun 10:474.

Frank SM, Forster L, Pawellek M, Malloni WM, Ahn S, Tse PU, Greenlee MW (2021) Visual attention modulates glutamate-glutamine levels in vestibular cortex: Evidence from magnetic resonance spectroscopy. The Journal of neuroscience : the official journal of the Society for Neuroscience.

Furlan AD, Pennick V, Bombardier C, van Tulder M, Group from the EB of the CBR (2009) 2009 Updated Method Guidelines for Systematic Reviews in the Cochrane Back Review Group. Spine 34:1929–1941.

Gao F, Edden RA, Li M, Puts NA, Wang G, Liu C, Zhao B, Wang H, Bai X, Zhao C (2013) Edited magnetic resonance spectroscopy detects an age-related decline in brain GABA levels. Neuroimage 78:75–82.

Gonçalves-Ribeiro J, Pina CC, Sebastião AM, Vaz SH (2019) Glutamate Transporters in Hippocampal LTD/LTP: Not Just Prevention of Excitotoxicity. Frontiers in Cellular Neuroscience 13:357.

Gordon EM et al. (2017) Precision Functional Mapping of Individual Human Brains. Neuron 95:791–807.e7.

Grames EM, Stillman AN, Tingley MW, Elphick CS (2019) An automated approach to identifying search terms for systematic reviews using keyword co-occurrence networks. Methods in Ecology and Evolution 10:1645–1654.

Gussew A, Rzanny R, Erdtel M, Scholle HC, Kaiser WA, Mentzel HJ, Reichenbach JR (2010) Time-resolved functional 1H MR spectroscopic detection of glutamate concentration changes in the brain during acute heat pain stimulation. NeuroImage 49:1895–1902.

Gutzeit A, Meier D, Froehlich JM, Hergan K, Kos S, V Weymarn C, Lutz K, Ettlin D, Binkert CA, Mutschler J, Sartoretti-Schefer S, Brügger M (2013) Differential NMR spectroscopy reactions of anterior/posterior and right/left insular subdivisions due to acute dental pain. European radiology 23:450–460.

Gutzeit A, Meier D, Meier ML, von Weymarn C, Ettlin DA, Graf N, Froehlich JM, Binkert CA, Brügger M (2011) Insula-specific responses induced by dental pain. A proton magnetic resonance spectroscopy study. European radiology 21:807–815.

Haddaway NR, Page MJ, Pritchard CC, McGuinness LA (2022) PRISMA2020: An R package and Shiny app for producing PRISMA 2020-compliant flow diagrams, with interactivity for optimised digital transparency and Open Synthesis. Campbell Systematic Reviews 18:e1230.

Hak T, Van Rhee H, Suurmond R (2016) How to interpret results of meta-analysis, (Version 1.0). Rotterdam, The Netherlands: Erasmus Rotterdam Institute of Management. Available at: www.erim.eur.nl/research-support/meta-essentials.

Harris AD, Saleh MG, Edden RAE (2017) Edited 1 H magnetic resonance spectroscopy in vivo: Methods and metabolites. Magn Reson Med 77:1377–1389.

Hasler G, van der Veen JW, Grillon C, Drevets WC, Shen J (2010) Effect of acute psychological stress on prefrontal GABA concentration determined by proton magnetic resonance spectroscopy. Am J Psychiatry 167:1226–1231.

Heba S, Puts NAJ, Kalisch T, Glaubitz B, Haag LM, Lenz M, Dinse HR, Edden RAE, Tegenthoff M, Schmidt-Wilcke T (2016) Local GABA Concentration Predicts Perceptual Improvements After Repetitive Sensory Stimulation in Humans. Cerebral Cortex 26:1295–1301.

Henning A (2018) Proton and multinuclear magnetic resonance spectroscopy in the human brain at ultra-high field strength: A review. NeuroImage 168:181–198.

Hermans L, Leunissen I, Pauwels L, Cuypers K, Peeters R, Puts NAJ, Edden RAE, Swinnen SP (2018) Brain GABA Levels Are Associated with Inhibitory Control Deficits in Older Adults. J Neurosci 38:7844–7851.

Hertz L, Rothman DL (2016) Glucose, Lactate, β-Hydroxybutyrate, Acetate, GABA, and Succinate as Substrates for Synthesis of Glutamate and GABA in the Glutamine-Glutamate/GABA Cycle. Adv Neurobiol 13:9–42.

Higgins JP (2011) Cochrane handbook for systematic reviews of interventions. Version 5.1. 0 [updated March 2011]. The Cochrane Collaboration. www.cochrane-handbook.org.

Higgins JPT, Altman DG, Gøtzsche PC, Jüni P, Moher D, Oxman AD, Savović J, Schulz KF, Weeks L, Sterne JAC (2011) The Cochrane Collaboration’s tool for assessing risk of bias in randomised trials. BMJ 343:d5928.

Higgins JPT, Thompson SG, Deeks JJ, Altman DG (2003) Measuring inconsistency in meta-analyses. BMJ 327:557–560.

Hoge RD, Atkinson J, Gill B, Crelier GR, Marrett S, Pike GB (1999) Stimulus-Dependent BOLD and Perfusion Dynamics in Human V1. NeuroImage 9:573–585.

Horder J, Petrinovic MM, Mendez MA, Bruns A, Takumi T, Spooren W, Barker GJ, Künnecke B, Murphy DG (2018) Glutamate and GABA in autism spectrum disorder—a translational magnetic resonance spectroscopy study in man and rodent models. Translational psychiatry 8:1–11.

Houtepen LC, Schür RR, Wijnen JP, Boer VO, Boks MPM, Kahn RS, Joëls M, Klomp DW, Vinkers CH (2017) Acute stress effects on GABA and glutamate levels in the prefrontal cortex: A 7T 1H magnetic resonance spectroscopy study. NeuroImage: Clinical 14:195–200.

Huang Z, Davis HHIV, Yue Q, Wiebking C, Duncan NW, Zhang J, Wagner N-F, Wolff A, Northoff G (2015) Increase in glutamate/glutamine concentration in the medial prefrontal cortex during mental imagery: A combined functional mrs and fMRI study. Human brain mapping 36:3204–3212.

Hui SCN et al. (2021) Frequency drift in MR spectroscopy at 3T. NeuroImage 241:118430.

IntHout J, Ioannidis JP, Borm GF (2014) The Hartung-Knapp-Sidik-Jonkman method for random effects meta-analysis is straightforward and considerably outperforms the standard DerSimonian-Laird method. BMC Medical Research Methodology 14:25.

Ioannidis JPA, Trikalinos TA (2007) The appropriateness of asymmetry tests for publication bias in meta-analyses: a large survey. CMAJ 176:1091–1096.

Ip IB, Bridge H (2021) Investigating the neurochemistry of the human visual system using magnetic resonance spectroscopy. Brain Struct Funct Available at: https://doi.org/10.1007/s00429-021-02273-0 [Accessed November 2, 2021].

Ip IB, Emir UE, Parker AJ, Campbell J, Bridge H (2019) Comparison of Neurochemical and BOLD Signal Contrast Response Functions in the Human Visual Cortex. J Neurosci 39:7968–7975.

Jahng G-H, Oh J, Lee D-W, Kim H-G, Rhee HY, Shin W, Paik J-W, Lee KM, Park S, Choe B-Y, Ryu C-W (2016) Glutamine and Glutamate Complex, as Measured by Functional Magnetic Resonance Spectroscopy, Alters During Face-Name Association Task in Patients with Mild Cognitive Impairment and Alzheimer’s Disease. Journal of Alzheimer’s Disease 52:145–159.

Jelen LA, King S, Horne CM, Lythgoe DJ, Young AH, Stone JM (2019) Functional magnetic resonance spectroscopy in patients with schizophrenia and bipolar affective disorder: Glutamate dynamics in the anterior cingulate cortex during a working memory task. European neuropsychopharmacology : the journal of the European College of Neuropsychopharmacology 29:222–234.

Jo S et al. (2014) GABA from reactive astrocytes impairs memory in mouse models of Alzheimer’s disease. Nat Med 20:886–896.

Juchem C, de Graaf RA (2017) B0 magnetic field homogeneity and shimming for in vivo magnetic resonance spectroscopy. Anal Biochem 529:17–29.

Just N, Faber C (2019) Probing activation-induced neurochemical changes using optogenetics combined with functional magnetic resonance spectroscopy: a feasibility study in the rat primary somatosensory cortex. Journal of Neurochemistry 150:402–419.

Just N, Sonnay S (2017) Investigating the Role of Glutamate and GABA in the Modulation of Transthalamic Activity: A Combined fMRI-fMRS Study. Front Physiol 8:30.

Just N, Xin L, Frenkel H, Gruetter R (2013) Characterization of sustained BOLD activation in the rat barrel cortex and neurochemical consequences. Neuroimage 74:343–351.

Kanowski M, Kaufmann J, Braun J, Bernarding J, Tempelmann C (2004) Quantitation of simulated short echo time 1H human brain spectra by LCModel and AMARES. Magnetic Resonance in Medicine 51:904–912.

Kiemes A, Davies C, Kempton MJ, Lukow PB, Bennallick C, Stone JM, Modinos G (2021) GABA, Glutamate and Neural Activity: A Systematic Review With Meta-Analysis of Multimodal 1H-MRS-fMRI Studies. Frontiers in Psychiatry 12:255.

Kim SY, Park JE, Lee YJ, Seo H-J, Sheen S-S, Hahn S, Jang B-H, Son H-J (2013a) Testing a tool for assessing the risk of bias for nonrandomized studies showed moderate reliability and promising validity. J Clin Epidemiol 66:408–414.

Kim T-H, Kang H-K, Jeong G-W (2013b) Assessment of brain metabolites change during visual sexual stimulation in healthy women using functional MR spectroscopy. The journal of sexual medicine 10:1001–1011.

Kim T-H, Kang H-K, Park K, Jeong G-W (2014) Localized brain metabolite changes during visual sexual stimulation in postmenopausal women: a pilot study using functional magnetic resonance spectroscopy. Menopause (New York, NY) 21:59–66.

Klose U (2008) Measurement sequences for single voxel proton MR spectroscopy. European journal of radiology 67:194–201.

Kolasinski J, Hinson EL, Zand APD, Rizov A, Emir UE, Stagg CJ (2019) The dynamics of cortical GABA in human motor learning. The Journal of Physiology 597:271–282.

Koush Y, de Graaf RA, Kupers R, Dricot L, Ptito M, Behar KL, Rothman DL, Hyder F (2021a) Metabolic underpinnings of activated and deactivated cortical areas in human brain. Journal of cerebral blood flow and metabolism : official journal of the International Society of Cerebral Blood Flow and Metabolism:271678X21989186–271678X21989186.

Koush Y, de Graaf RA, Kupers R, Dricot L, Ptito M, Behar KL, Rothman DL, Hyder F (2021b) Metabolic underpinnings of activated and deactivated cortical areas in human brain. J Cereb Blood Flow Metab 41:986–1000.

Krüger G, Kleinschmidt A, Frahm J (1998) Stimulus dependence of oxygenation-sensitive MRI responses to sustained visual activation. NMR Biomed 11:75–79.

Kühn S, Schubert F, Mekle R, Wenger E, Ittermann B, Lindenberger U, Gallinat J (2016) Neurotransmitter changes during interference task in anterior cingulate cortex: evidence from fMRI-guided functional MRS at 3 T. Brain structure & function 221:2541–2551.

Kurcyus K, Annac E, Hanning NM, Harris AD, Oeltzschner G, Edden R, Riedl V (2018) Opposite Dynamics of GABA and Glutamate Levels in the Occipital Cortex during Visual Processing. The Journal of neuroscience : the official journal of the Society for Neuroscience 38:9967–9976.

Kuwabara T, Watanabe H, Tsuji S, Yuasa T (1995) Lactate rise in the basal ganglia accompanying finger movements: a localized1H-MRS study. Brain Research 670:326–328.

Lajeunesse MJ (2016) Facilitating systematic reviews, data extraction and meta-analysis with the metagear package for r. Methods in Ecology and Evolution 7:323–330.

Lally N, Mullins PG, Roberts MV, Price D, Gruber T, Haenschel C (2014) Glutamatergic correlates of gamma-band oscillatory activity during cognition: a concurrent ER-MRS and EEG study. NeuroImage 85 Pt 2:823–833.

Li C-T, Yang K-C, Lin W-C (2019) Glutamatergic Dysfunction and Glutamatergic Compounds for Major Psychiatric Disorders: Evidence From Clinical Neuroimaging Studies. Frontiers in Psychiatry 9 Available at: https://www.frontiersin.org/articles/10.3389/fpsyt.2018.00767 [Accessed July 20, 2022].

Ligneul C, Fernandes FF, Shemesh N (2021) High temporal resolution functional magnetic resonance spectroscopy in the mouse upon visual stimulation. NeuroImage 234:117973.

Lin A et al. (2021) Minimum Reporting Standards for in vivo Magnetic Resonance Spectroscopy (MRSinMRS): Experts’ consensus recommendations. NMR Biomed 34:e4484.

Lin L, Chu H (2018) Quantifying Publication Bias in Meta-Analysis. Biometrics 74:785–794.

Lin Y, Stephenson MC, Xin L, Napolitano A, Morris PG (2012) Investigating the metabolic changes due to visual stimulation using functional proton magnetic resonance spectroscopy at 7 T. Journal of cerebral blood flow and metabolism : official journal of the International Society of Cerebral Blood Flow and Metabolism 32:1484–1495.

Lynn J, Woodcock EA, Anand C, Khatib D, Stanley JA (2018a) Differences in steady-state glutamate levels and variability between “non-task-active” conditions: Evidence from (1)H fMRS of the prefrontal cortex. NeuroImage 172:554–561.

Lynn J, Woodcock EA, Anand C, Khatib D, Stanley JA (2018b) Differences in steady-state glutamate levels and variability between ‘non-task-active’ conditions: Evidence from 1H fMRS of the prefrontal cortex. Neuroimage 172:554–561.

Ma L-L, Wang Y-Y, Yang Z-H, Huang D, Weng H, Zeng X-T (2020) Methodological quality (risk of bias) assessment tools for primary and secondary medical studies: what are they and which is better? Military Medical Research 7:7.

Maddock RJ, Casazza GA, Buonocore MH, Tanase C (2011) Vigorous exercise increases brain lactate and Glx (glutamate+glutamine): a dynamic 1H-MRS study. NeuroImage 57:1324–1330.

Magalhães R, Novais A, Barrière DA, Marques P, Marques F, Sousa JC, Cerqueira JJ, Cachia A, Jay TM, Bottlaender M, Sousa N, Mériaux S, Boumezbeur F (2019) A Resting-State Functional MR Imaging and Spectroscopy Study of the Dorsal Hippocampus in the Chronic Unpredictable Stress Rat Model. J Neurosci 39:3640–3650.

Magistretti PJ, Allaman I (2015) A Cellular Perspective on Brain Energy Metabolism and Functional Imaging. Neuron 86:883–901.

Mangia S, Giove F, DiNuzzo M (2012) Metabolic Pathways and Activity-Dependent Modulation of Glutamate Concentration in the Human Brain. Neurochem Res 37:2554–2561.

Mangia S, Giove F, Tkáč I, Logothetis NK, Henry P-G, Olman CA, Maraviglia B, Di Salle F, Uğurbil K (2009) Metabolic and Hemodynamic Events after Changes in Neuronal Activity: Current Hypotheses, Theoretical Predictions and in vivo NMR Experimental Findings. J Cereb Blood Flow Metab 29:441–463.

Mangia S, Tkác I, Gruetter R, Van de Moortele P-F, Maraviglia B, Uğurbil K (2007) Sustained neuronal activation raises oxidative metabolism to a new steady-state level: evidence from 1H NMR spectroscopy in the human visual cortex. Journal of cerebral blood flow and metabolism : official journal of the International Society of Cerebral Blood Flow and Metabolism 27:1055–1063.

Marenco S, Savostyanova AA, van der Veen JW, Geramita M, Stern A, Barnett AS, Kolachana B, Radulescu E, Zhang F, Callicott JH, Straub RE, Shen J, Weinberger DR (2010) Genetic Modulation of GABA Levels in the Anterior Cingulate Cortex by GAD1 and COMT. Neuropsychopharmacol 35:1708–1717.

Marjańska M, Deelchand DK, Kreis R, Team the 2016 IMSGFC (2022) Results and interpretation of a fitting challenge for MR spectroscopy set up by the MRS study group of ISMRM. Magnetic Resonance in Medicine 87:11–32.

Martínez-Maestro M, Labadie C, Möller HE (2019) Dynamic metabolic changes in human visual cortex in regions with positive and negative blood oxygenation level-dependent response. Journal of cerebral blood flow and metabolism : official journal of the International Society of Cerebral Blood Flow and Metabolism 39:2295–2307.

Mekle R, Kühn S, Pfeiffer H, Aydin S, Schubert F, Ittermann B (2017) Detection of metabolite changes in response to a varying visual stimulation paradigm using short-TE (1) H MRS at 7 T. NMR in biomedicine 30.

Mescher M, Merkle H, Kirsch J, Garwood M, Gruetter R (1998) Simultaneous in vivo spectral editing and water suppression. NMR Biomed 11:266–272.

Michels L, Martin E, Klaver P, Edden R, Zelaya F, Lythgoe DJ, Lüchinger R, Brandeis D, O’Gorman RL (2012) Frontal GABA Levels Change during Working Memory. PLOS ONE 7:e31933.

Mikkelsen M, Loo RS, Puts NAJ, Edden RAE, Harris AD (2018) Designing GABA-Edited Magnetic Resonance Spectroscopy Studies: Considerations of Scan Duration, Signal-To-Noise Ratio and Sample Size. J Neurosci Methods 303:86–94.

Minati L, Aquino D, Bruzzone MG, Erbetta A (2010) Quantitation of normal metabolite concentrations in six brain regions by in-vivo 1H-MR spectroscopy. J Med Phys 35:154–163.

Mullins PG (2018) Towards a theory of functional magnetic resonance spectroscopy (fMRS): A meta-analysis and discussion of using MRS to measure changes in neurotransmitters in real time. Scand J Psychol 59:91–103.

Mullins PG, McGonigle DJ, O’Gorman RL, Puts NAJ, Vidyasagar R, Evans CJ, Cardiff Symposium on MRS of GABA, Edden RAE (2014) Current practice in the use of MEGA-PRESS spectroscopy for the detection of GABA. Neuroimage 86:43–52.

Mullins PG, Rowland LM, Jung RE, Sibbitt WL (2005) A novel technique to study the brain’s response to pain: Proton magnetic resonance spectroscopy. NeuroImage 26:642–646.

Murphy KR, Aguinis H (2019) HARKing: How Badly Can Cherry-Picking and Question Trolling Produce Bias in Published Results? Journal of Business and Psychology 34:1–17.

Muthukumaraswamy SD, Evans CJ, Edden RAE, Wise RG, Singh KD (2012) Individual variability in the shape and amplitude of the BOLD-HRF correlates with endogenous GABAergic inhibition. Human Brain Mapping 33:455–465.

Nakagawa S, Lagisz M, Jennions MD, Koricheva J, Noble DW, Parker TH, Sánchez-Tójar A, Yang Y, O’Dea RE (2021) Methods for testing publication bias in ecological and evolutionary meta-analyses. Methods in Ecology and Evolution.

Nakahara T et al. (2022) Glutamatergic and GABAergic metabolite levels in schizophrenia-spectrum disorders: a meta-analysis of 1H-magnetic resonance spectroscopy studies. Mol Psychiatry 27:744–757.

Narayanan NS, Laubach M (2017) Inhibitory Control: Mapping Medial Frontal Cortex. Current Biology 27:R148–R150.

Near J, Simpson R, Cowen P, Jezzard P (2011) Efficient γ-aminobutyric acid editing at 3T without macromolecule contamination: MEGA-SPECIAL. NMR Biomed 24:1277–1285.

Öz G, Tkáč I (2011) Short-echo, single-shot, full-intensity proton magnetic resonance spectroscopy for neurochemical profiling at 4 T: Validation in the cerebellum and brainstem. Magnetic Resonance in Medicine 65:901–910.

Page MJ et al. (2021) The PRISMA 2020 statement: an updated guideline for reporting systematic reviews. BMJ 372:n71.

Paredes RG, Agmo A (1992) GABA and behavior: the role of receptor subtypes. Neurosci Biobehav Rev 16:145–170.

Peek AL, Rebbeck T, Puts NA, Watson J, Aguila M-ER, Leaver AM (2020) Brain GABA and glutamate levels across pain conditions: A systematic literature review and meta-analysis of 1H-MRS studies using the MRS-Q quality assessment tool. Neuroimage 210:116532.

Porges EC, Woods AJ, Lamb DG, Williamson JB, Cohen RA, Edden RAE, Harris AD (2017) Impact of tissue correction strategy on GABA-edited MRS findings. NeuroImage 162:249–256.

Pouwels PJ, Frahm J (1998) Regional metabolite concentrations in human brain as determined by quantitative localized proton MRS. Magn Reson Med 39:53–60.

Prichard JW (1992) Magnetic resonance spectroscopy of the brain. Clin Chim Acta 206:115–123.

Pustejovsky JE, Tipton E (2021) Meta-analysis with Robust Variance Estimation: Expanding the Range of Working Models. Prev Sci.

Puts NAJ, Edden RAE (2012) In vivo magnetic resonance spectroscopy of GABA: A methodological review. Prog Nucl Magn Reson Spectrosc 60:29–41.

Puts NAJ, Edden RAE, Evans CJ, McGlone F, McGonigle DJ (2011) Regionally Specific Human GABA Concentration Correlates with Tactile Discrimination Thresholds. J Neurosci 31:16556–16560.

Rae C, Nasrallah FA, Griffin JL, Balcar VJ (2009) Now I know my ABC. A systems neurochemistry and functional metabolomic approach to understanding the GABAergic system. J Neurochem 109 Suppl 1:109–116.

Rae CD (2014) A Guide to the Metabolic Pathways and Function of Metabolites Observed in Human Brain 1H Magnetic Resonance Spectra. Neurochem Res 39:1–36.

Rideaux R (2020) Temporal Dynamics of GABA and Glx in the Visual Cortex. eNeuro 7 Available at: https://www.eneuro.org/content/7/4/ENEURO.0082-20.2020 [Accessed January 7, 2022].

Rideaux R (2021) No balance between glutamate+glutamine and GABA+ in visual or motor cortices of the human brain: A magnetic resonance spectroscopy study. NeuroImage 237:118191.

Rosenthal R (1979) The file drawer problem and tolerance for null results. Psychological Bulletin 86:638–641.

Rosenthal R (1986) Meta-Analytic Procedures for Social Science Research Sage Publications: Beverly Hills, 1984, 148 pp. Educational Researcher 15:18–20.

Rothman DL, Behar KL, Hyder F, Shulman RG (2003) In vivo NMR studies of the glutamate neurotransmitter flux and neuroenergetics: implications for brain function. Annu Rev Physiol 65:401–427.

Ruppert D, Wand MP (1994) Multivariate Locally Weighted Least Squares Regression. The Annals of Statistics 22:1346–1370.

Sanaei Nezhad F, Anton A, Michou E, Jung J, Parkes LM, Williams SR (2018) Quantification of GABA, glutamate and glutamine in a single measurement at 3 T using GABA-edited MEGA-PRESS. NMR in Biomedicine 31:e3847.

Schaller B, Mekle R, Xin L, Kunz N, Gruetter R (2013) Net increase of lactate and glutamate concentration in activated human visual cortex detected with magnetic resonance spectroscopy at 7 tesla. Journal of neuroscience research 91:1076–1083.

Schaller B, Xin L, O’Brien K, Magill AW, Gruetter R (2014) Are glutamate and lactate increases ubiquitous to physiological activation? A (1)H functional MR spectroscopy study during motor activation in human brain at 7Tesla. NeuroImage 93 Pt 1:138–145.

Schür RR, Draisma LWR, Wijnen JP, Boks MP, Koevoets MGJC, Joëls M, Klomp DW, Kahn RS, Vinkers CH (2016) Brain GABA levels across psychiatric disorders: A systematic literature review and meta-analysis of 1H-MRS studies. Human Brain Mapping 37:3337–3352.

Shi L, Lin L (2019) The trim-and-fill method for publication bias: practical guidelines and recommendations based on a large database of meta-analyses. Medicine 98:e15987.

Sibson NR, Dhankhar A, Mason GF, Rothman DL, Behar KL, Shulman RG (1998) Stoichiometric coupling of brain glucose metabolism and glutamatergic neuronal activity. PNAS 95:316–321.

Siniatchkin M, Sendacki M, Moeller F, Wolff S, Jansen O, Siebner H, Stephani U (2012) Abnormal changes of synaptic excitability in migraine with aura. Cerebral cortex (New York, NY : 1991) 22:2207–2216.

Stagg CJ, Bachtiar V, Johansen-Berg H (2011) The Role of GABA in Human Motor Learning. Curr Biol 21:480–484.

Stagg CJ, Best JG, Stephenson MC, O’Shea J, Wylezinska M, Kincses ZT, Morris PG, Matthews PM, Johansen-Berg H (2009) Polarity-Sensitive Modulation of Cortical Neurotransmitters by Transcranial Stimulation. J Neurosci 29:5202–5206.

Stanley JA, Burgess A, Khatib D, Ramaseshan K, Arshad M, Wu H, Diwadkar VA (2017) Functional dynamics of hippocampal glutamate during associative learning assessed with in vivo (1)H functional magnetic resonance spectroscopy. NeuroImage 153:189–197.

Stanley JA, Raz N (2018) Functional Magnetic Resonance Spectroscopy: The “New” MRS for Cognitive Neuroscience and Psychiatry Research. Frontiers in Psychiatry 9:76.

Steel A, Mikkelsen M, Edden RA, Robertson CE (2020) Regional balance between glutamate+ glutamine and GABA+ in the resting human brain. Neuroimage 220:117112.

Steen RG, Hamer RM, Lieberman JA (2005) Measurement of brain metabolites by 1H magnetic resonance spectroscopy in patients with schizophrenia: a systematic review and meta-analysis. Neuropsychopharmacology 30:1949–1962.

Sterne JAC, Egger M (2001) Funnel plots for detecting bias in meta-analysis: Guidelines on choice of axis. Journal of Clinical Epidemiology 54:1046–1055.

Suurmond R, van Rhee H, Hak T (2017) Introduction, comparison, and validation of Meta-Essentials: A free and simple tool for meta-analysis. Research Synthesis Methods 8:537–553.

Takado Y, Takuwa H, Sampei K, Urushihata T, Takahashi M, Shimojo M, Uchida S, Nitta N, Shibata S, Nagashima K, Ochi Y, Ono M, Maeda J, Tomita Y, Sahara N, Near J, Aoki I, Shibata K, Higuchi M (2021) MRS-measured glutamate versus GABA reflects excitatory versus inhibitory neural activities in awake mice. J Cereb Blood Flow Metab:0271678X211045449.

Tang X, Jaenisch R, Sur M (2021) The role of GABAergic signalling in neurodevelopmental disorders. Nat Rev Neurosci 22:290–307.

Taylor R, Neufeld RWJ, Schaefer B, Densmore M, Rajakumar N, Osuch EA, Williamson PC, Théberge J (2015a) Functional magnetic resonance spectroscopy of glutamate in schizophrenia and major depressive disorder: anterior cingulate activity during a color-word Stroop task. NPJ schizophrenia 1:15028–15028.

Taylor R, Schaefer B, Densmore M, Neufeld RWJ, Rajakumar N, Williamson PC, Théberge J (2015b) Increased glutamate levels observed upon functional activation in the anterior cingulate cortex using the Stroop Task and functional spectroscopy. Neuroreport 26:107–112.

Terpstra M, Cheong I, Lyu T, Deelchand DK, Emir UE, Bednařík P, Eberly LE, Öz G (2016) Test-retest reproducibility of neurochemical profiles with short-echo, single-voxel MR spectroscopy at 3T and 7T. Magn Reson Med 76:1083–1091.

Terrin N, Schmid CH, Lau J, Olkin I (2003) Adjusting for publication bias in the presence of heterogeneity. Stat Med 22:2113–2126.

Tricco AC, Lillie E, Zarin W, O’Brien KK, Colquhoun H, Levac D, Moher D, Peters MD, Horsley T, Weeks L (2018) PRISMA extension for scoping reviews (PRISMA-ScR): checklist and explanation. Annals of internal medicine 169:467–473.

Valtcheva S, Venance L (2019) Control of Long-Term Plasticity by Glutamate Transporters. Frontiers in Synaptic Neuroscience 11:10.

Viechtbauer W (2010) Conducting meta-analyses in R with the metafor package. Journal of statistical software 36:1–48.

Vijayakumari AA, Menon RN, Thomas B, Arun TM, Nandini M, Kesavadas C (2020) Glutamatergic response to a low load working memory paradigm in the left dorsolateral prefrontal cortex in patients with mild cognitive impairment: a functional magnetic resonance spectroscopy study. Brain imaging and behavior 14:451–459.

Vijayakumari AA, Thomas B, Menon RN, Kesavadas C (2018) Association between glutamate/glutamine and blood oxygen level dependent signal in the left dorsolateral prefrontal region during verbal working memory. Neuroreport 29:478–482.

Volovyk O, Tal A (2020) Increased Glutamate concentrations during prolonged motor activation as measured using functional Magnetic Resonance Spectroscopy at 3T. NeuroImage 223:117338– 117338.

Wilson M et al. (2019) Methodological consensus on clinical proton MRS of the brain: Review and recommendations. Magnetic Resonance in Medicine 82:527–550.

Woodcock EA, Anand C, Khatib D, Diwadkar VA, Stanley JA (2018) Working Memory Modulates Glutamate Levels in the Dorsolateral Prefrontal Cortex during (1)H fMRS. Frontiers in psychiatry 9:66–66.

Woodcock EA, Greenwald MK, Khatib D, Diwadkar VA, Stanley JA (2019) Pharmacological stress impairs working memory performance and attenuates dorsolateral prefrontal cortex glutamate modulation. NeuroImage 186:437–445.

Yashiro K, Corlew R, Philpot BD (2005) Visual Deprivation Modifies Both Presynaptic Glutamate Release and the Composition of Perisynaptic/Extrasynaptic NMDA Receptors in Adult Visual Cortex. J Neurosci 25:11684–11692.

Yizhar O, Fenno LE, Prigge M, Schneider F, Davidson TJ, O’Shea DJ, Sohal VS, Goshen I, Finkelstein J, Paz JT, Stehfest K, Fudim R, Ramakrishnan C, Huguenard JR, Hegemann P, Deisseroth K (2011) Neocortical excitation/inhibition balance in information processing and social dysfunction. Nature 477:171–178.

Zhang X, Liang M, Qin W, Wan B, Yu C, Ming D (2020) Gender Differences Are Encoded Differently in the Structure and Function of the Human Brain Revealed by Multimodal MRI. Frontiers in Human Neuroscience 14 Available at: https://www.frontiersin.org/article/10.3389/fnhum.2020.00244 [Accessed January 25, 2022].

Zhu X-H, Chen W (2001) Observed BOLD effects on cerebral metabolite resonances in human visual cortex during visual stimulation: A functional 1H MRS study at 4 T. Magnetic Resonance in Medicine 46:841–847.

Zöllner HJ, Považan M, Hui SCN, Tapper S, Edden RAE, Oeltzschner G (2021) Comparison of different linear-combination modeling algorithms for short-TE proton spectra. NMR in Biomedicine 34:e4482.

Zöllner HJ, Tapper S, Hui SCN, Barker PB, Edden RAE, Oeltzschner G (2022) Comparison of linear combination modeling strategies for edited magnetic resonance spectroscopy at 3 T. NMR Biomed 35:e4618.

