## Supplemental Information for "Functional MRS studies of GABA and Glutamate/Glx – a systematic review and meta-analysis"

Supplementary table 1.

Search strategy from OVID

| **Mesh** | **Keyword** |
| --- | --- |
| Magnetic Resonance spectroscopy | spectroscop* |
| Proton Magnetic resonance spectroscopy | magnetic resonance and (imag* or spectroscop*) |
|  | magnetic resonance spectroscopy |
|  | proton magnetic resonance spectroscopy |
|  | mega press |
|  | 1hmrs |
|  | 1h-mrs |
|  | in-vivo mrs |
|  | Mr |
|  | Mrs |
|  | mr-spectro* |
|  | Nmr |
|  | in-vivo nmr |
|  | functional magnetic resonance spectroscopy.mp. |
|  | fmrs.mp. |
|  | functional mrs.mp. |
| AND | |
| Functional Status | functional.mp. |
|  | dynamic.mp. |
|  | func*.mp. |
| AND | |
| gamma-Aminobutyric Acid | Neurotransmitter.mp |
| Glutamic Acid | neurochemical.mp |
| Glutamine |  |
| Brain Chemistry |  |
| Neurotransmitter Agents |  |

Supplementary table 2. results from Egger’s regression test

| Data | Metabolite | Adjusted estimate effect size | 95%CI | p |
| --- | --- | --- | --- | --- |
| Mean | Glu/Glx | -0.791 | -1.21, -0.37 | <0.001 |
|  | Glu | 0.857 | -1.37, -0.35 | <0.001 |
|  | Glx | -0.478 | -1.01, 0.05 | 0.034 |
|  | GABA | -0.881 | -2.21, 0.45 | 0.302 |
| %Change | Glu/Glx | -1.607 | -2.16, -1.05 | <0.001 |
|  | Glu | -1.707 | -2.35, -1.07 | <0.001 |
|  | Glx | -0.863 | -2.09, 0.36 | 0.092 |
|  | GABA | -0.198 | -1.70, 1.31 | 0.887 |

Supplementary table 3. results from trim-and-fill test. The effect sizes reported here in Table X is also the estimate effect sizes of aggregates, since no study was added to the analysis via trim-and-fill.

| Data type | Metabolite | Estimate effect size | 95%CI | zval | p | tau^2^ | I^2^ |
| --- | --- | --- | --- | --- | --- | --- | --- |
| Mean | Glu/Glx | 0.09 | -0.09, 0.28 | 1.07 | 0.29 | 0.16 | 85.56 |
|  | Glu | 0.21 | -0.04, 0.46 | 1.75 | 0.10 | 0.20 | 85.77 |
|  | Glx | 0.07 | -0.13, 0.27 | 0.76 | 0.46 | 0.08 | 76.63 |
|  | GABA | -0.26 | -0.61, 0.09 | -1.67 | 0.13 | 0.24 | 89.00 |
| %Change | Glu/Glx | 0.12 | -0.20, 0.45 | 0.81 | 0.43 | 0.36 | 91.31 |
|  | Glu | 0.11 | -0.35, 0.57 | 0.51 | 0.62 | 0.49 | 93.14 |
|  | Glx | 0.14 | -0.13, 0.42 | 1.19 | 0.27 | 0.06 | 62.79 |
|  | GABA | -0.10 | -0.33, 0.13 | -1.02 | 0.33 | 0.07 | 68.93 |

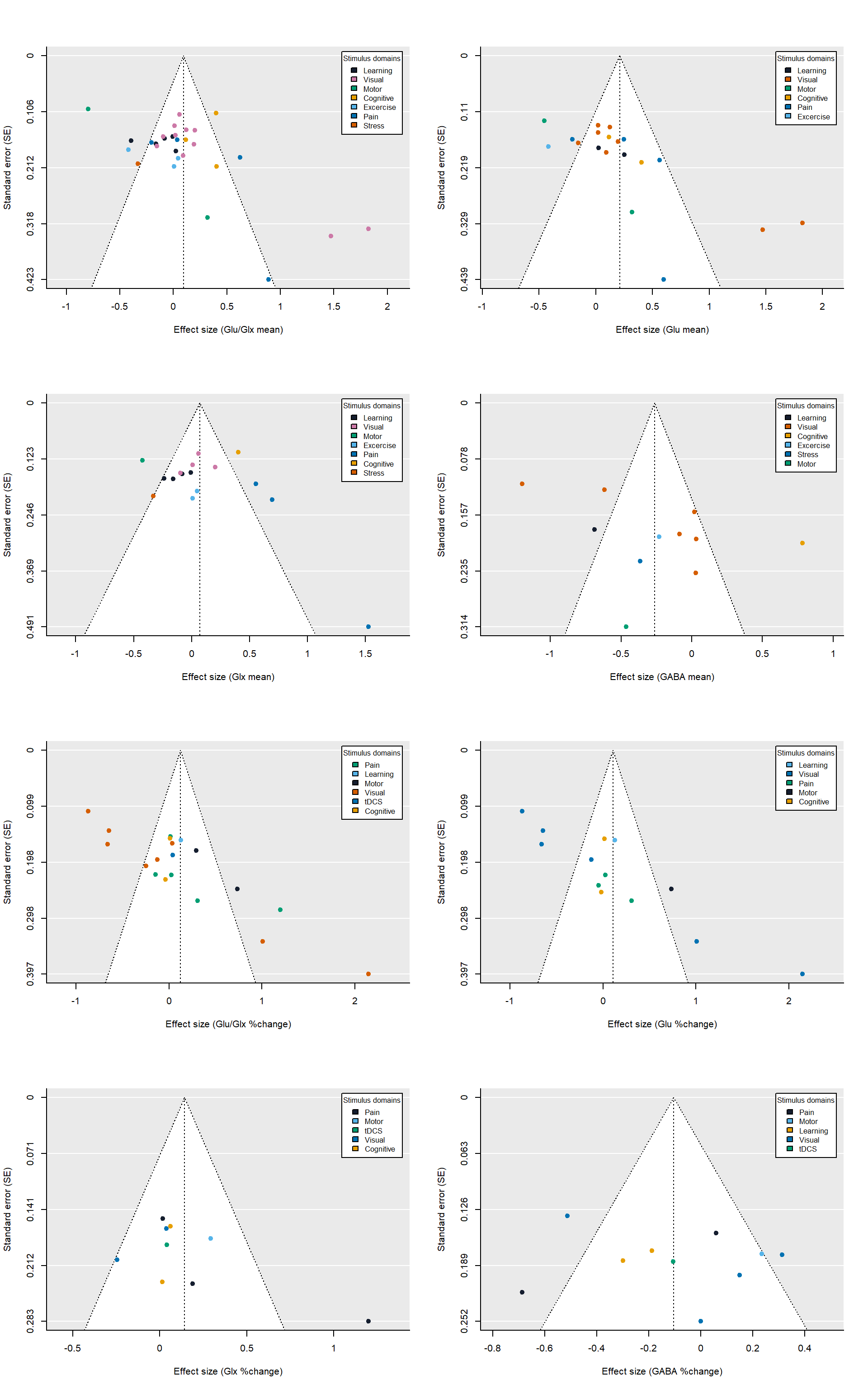

Supplementary figure 1. Funnel plot color-coded with stimulus domains

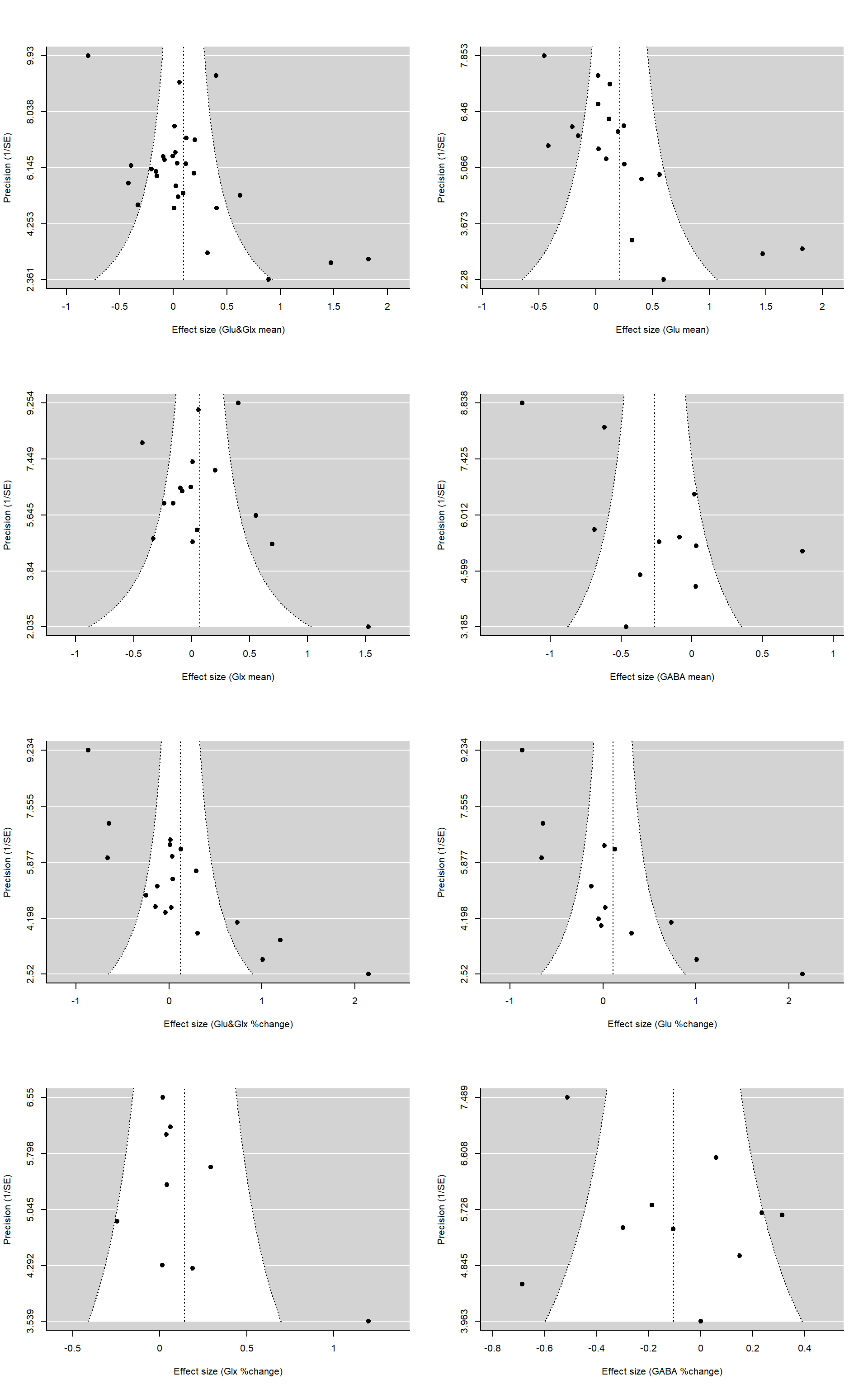

Supplementary figure 2. funnel plots from trim-and-fill test

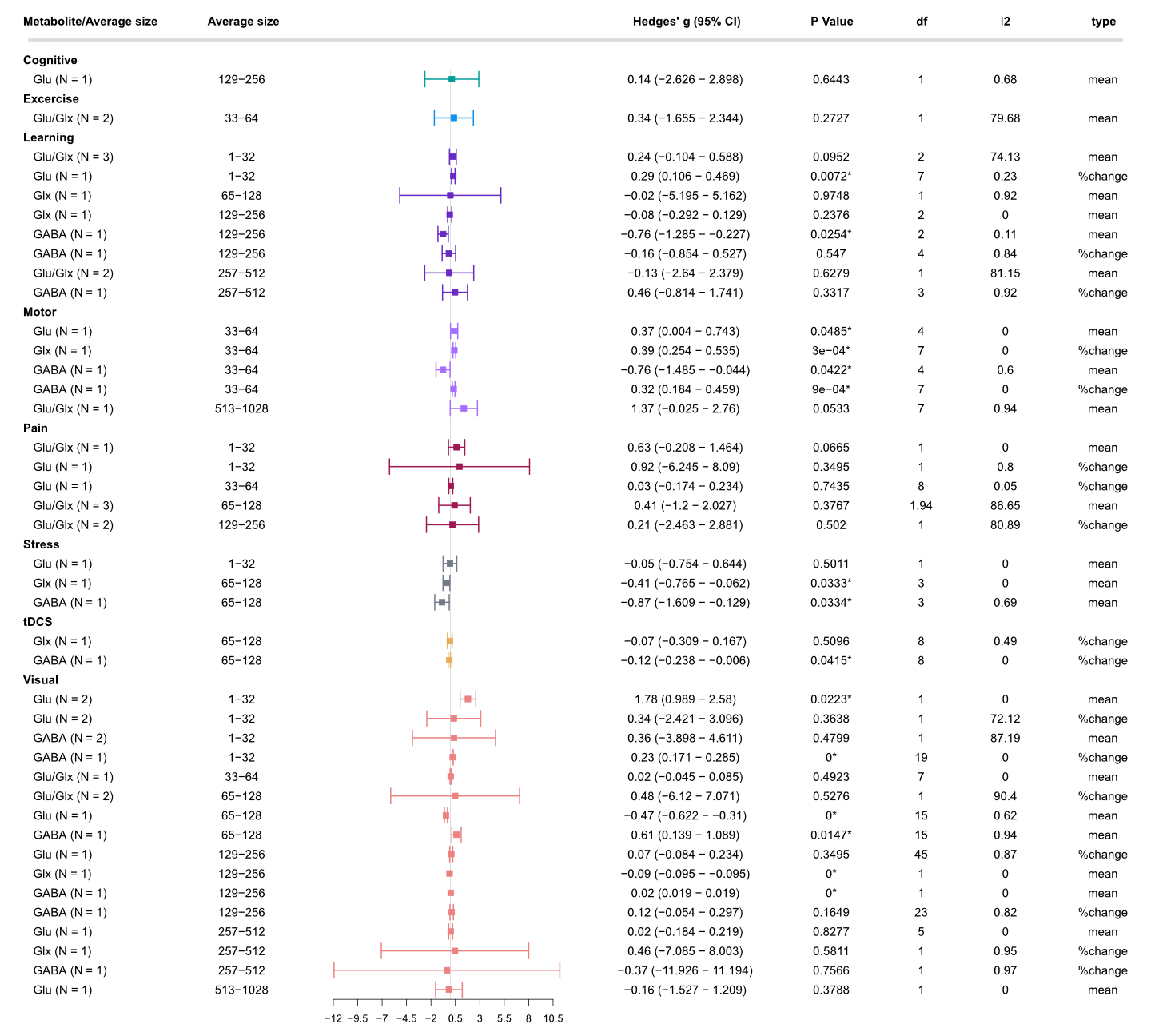

Supplementary Figure 3: Influence of range of average size on metabolite levels based on type of stimulus domains. N: number of studies included; I^2^: I^2^ index for heterogeneity. *Statistically significant at p <0.05, and at p <0.01 when the degrees of freedom < 4 for RVE t-tests.
